# Integrative and comparative genomic analyses of mammalian macrophage responses to intracellular mycobacterial pathogens

**DOI:** 10.1101/2023.07.14.549042

**Authors:** Thomas J. Hall, Gillian P. McHugo, Michael P. Mullen, James A. Ward, Kate E. Killick, John A. Browne, Stephen V. Gordon, David E. MacHugh

## Abstract

*Mycobacterium tuberculosis*, the causative agent of human tuberculosis (hTB), is currently classed as the thirteenth leading cause of death worldwide. *Mycobacterium bovis*, a close evolutionary relative of *M. tuberculosis*, causes bovine tuberculosis (bTB) and is one of the most damaging infectious diseases to livestock agriculture. Previous studies have shown that the pathogenesis of bTB disease is comparable to hTB disease, and that the bovine and human alveolar macrophage (bAM and hAM, respectively) transcriptomes are extensively reprogrammed in response to infection with these intracellular mycobacterial pathogens. However, although *M. bovis* and *M. tuberculosis* share over 99% identity at the genome level, the innate immune responses to these pathogens have been shown to be different in human or cattle hosts.

In this study, a multi-omics integrative approach was applied to encompass functional genomics and GWAS data sets across the two primary hosts (*Bos taurus* and *Homo sapiens*) and both pathogens (*M. bovis* and *M. tuberculosis*). Four different experimental infection groups were used, each with parallel non-infected control cells: 1) bAM infected with *M. bovis*, 2) bAM infected with *M. tuberculosis*, 3) hAM infected with *M. tuberculosis*, and 4) human monocyte-derived macrophages (hMDM) infected with *M. tuberculosis*. RNA-seq data from these experiments 24 hours post-infection (24 hpi) was analysed using three separate computational pipelines: 1) differentially expressed genes, 2) differential gene expression interaction networks, and 3) combined pathway analysis. The results of these analyses were then integrated with high-resolution bovine and human GWAS data sets to detect novel quantitative trait loci (QTLs) for resistance to mycobacterial infection and resilience to disease. Results from this study revealed common and unique response macrophage pathways for both pathogens and identified 32 genes (12 bovine and 20 human) significantly enriched for SNPs associated with disease resistance, the majority of which encode key components of the NF-κB signalling pathway and that also drive formation of the granuloma.

## 1. Introduction

The phylum *Actinobacteria* represents one of the largest taxonomic groups among the 18 major lineages currently recognized within the domain Bacteria, including five subclasses and 14 suborders. It comprises gram-positive bacteria with a high genomic DNA GC content, ranging from 51% in some *Corynebacterium* to more than 70% in *Streptomyces* and *Frankia* (Ventura *et al*. 2007). Two important disease-causing species of Actinobacteria, *Mycobacterium tuberculosis* and *Mycobacterium bovis*, are members of the *M. tuberculosis* complex (MTBC) and are characterised by 99.95% similarity at the nucleotide level (Garnier *et al*. 2003). Both species cause tuberculosis (TB) in mammals, with *M. tuberculosis* displaying a specific host preference for humans, with this MTBC strain not environmentally maintained by animal reservoirs (Brites & Gagneux 2015; Gagneux 2018).

*M. tuberculosis* is one of the most deadly human pathogens, and until the COVID-19 pandemic was the leading cause of death from a single infectious agent, ranking above HIV/AIDS with approximately 1.6 million deaths in 2021 and is the thirteenth leading cause of death overall (World Health Organization 2022). In 2021, there were also approximately 450,000 new cases that exhibited resistance to rifampicin (RR-TB), the most effective first-line drug, or that were multidrug-resistant (MDR-TB). Most deaths from hTB could be prevented with early diagnosis and appropriate treatment.

*M. bovis*, the causative agent of bovine tuberculosis (bTB), is considered to be one of the most damaging infectious diseases to the global agricultural industry, with the global costs of disease conservatively estimated to cost €3 billion annually, imposing huge economic losses on farmers of infected herds (Steele 1995; Schiller *et al*. 2010; Waters *et al*. 2012). *Mycobacterium bovis* can cause zoonotic TB (zTB) with serious implications for human health (Olea-Popelka *et al*. 2017; Vayr *et al*. 2018; Kock *et al*. 2021). Due to the highly infectious nature of the pathogen, early detection and removal of infected animals is the most effective control measure (Clegg *et al*. 2011). In contrast to *M. tuberculosis*, *M. bovis* has a wide host range and can infect a broad spectrum of domestic and wild mammals including taurine and zebu cattle (*Bos taurus* and *B. indicus*, respectively), sheep (*Ovis aries*), goats (*Capra hircus*), llamas and alpacas (*Lama* spp.), pigs and wild boar (*Sus scrofa*), European badgers (*Meles meles*), brushtail possums (*Trichosurus vulpecula*), various species of deer (Cervidae), African and Asian elephants (*Loxodonta* spp. and *Elephas maximus*), African buffalo (*Syncerus caffer*), and many Felidae species including African lions (*Panthera leo*), leopards (*Panthera pardus*), and cheetahs (*Acinonyx jubatus*) (Pesciaroli *et al*. 2014; Malone & Gordon 2017; Gormley & Corner 2018).

Considered together, *M. tuberculosis* and *M. bovis* represent enormous burdens on global health systems and animal agriculture worldwide. Previous studies have shown that the pathogenesis of bTB disease in cattle is comparable to hTB disease and many aspects of *M. bovis* infection are also characteristic of *M. tuberculosis* infection (Neill *et al*. 2001; Russell 2003; Cassidy 2006; Pollock *et al*. 2006; Waters *et al*. 2014). Consequently, *M. bovis* infection of cattle and bTB disease are now recognised as a valuable model for understanding hTB caused by *M. tuberculosis* (Van Rhijn *et al*. 2008; Waters *et al*. 2011; Waters & Palmer 2015; Williams & Orme 2016; Gong *et al*. 2020). Transmission of both MTBC strains is via inhalation of contaminated aerosol droplets and the primary site of infection is the lungs (Weiss & Schaible 2015; Dorhoi & Kaufmann 2016), where these pathogens encounter resident alveolar macrophages (AM), the host’s first line of defence, which normally phagocytise and destroy airborne bacteria (O’Garra *et al*. 2013; Cliff *et al*. 2015). However, MTBC mycobacteria such as *M. bovis* and *M. tuberculosis* can persist and replicate within alveolar macrophages via a plethora of evolved evasion mechanisms that subvert and interfere with host immune responses (Cambier *et al*. 2014; Schorey & Schlesinger 2016; Awuh & Flo 2017; Chandra *et al*. 2022). Some of these mechanisms encompass the early secretory antigenic target-6 (ESAT-6) secretion system-1 (ESX-1), which facilitates escape from the macrophage phagosome into the cytosol where it replicates (Goldberg *et al*. 2014). The phagosomes that form when macrophages digest an invading microorganism normally go through rapid fusion with lysosomes; however, this is not the case with AM that phagocytize the pathogenic MTBC strains (Hmama *et al*. 2015).

Previous studies, including those performed by our research group, have demonstrated that the bovine and human alveolar macrophage (bAM and hAM) transcriptomes are extensively reprogrammed in response to infection with *M. bovis* and *M. tuberculosis* (Nalpas *et al*. 2015; Vegh *et al*. 2015; Lavalett *et al*. 2017; Malone *et al*. 2018; Papp *et al*. 2018; Hall *et al*. 2020; Hall *et al*. 2021; Mendonca *et al*. 2023). These studies have also highlighted a complex system of gene expression regulation that drives host-pathogen interactions and innate immune response pathway execution, which are functionally associated with many macrophage processes that control or eliminate intracellular microbes. What remains unclear, however, is the innate immune response genes and pathways that are species-specific or common to both bAM and hAM infected with *M. bovis* and *M. tuberculosis*, respectively.

Even though *M. bovis* and *M. tuberculosis* share 99.95% identity at the genome level, the innate immune responses to the pathogens can be characteristically different. Though the two pathogens encounter common immune response pathways in their preferred host, there are certain genes that are activated or repressed in response to a specific MTBC pathogen. For example, a study by our group investigated the responses of bAM to *M. bovis* and *M. tuberculosis* at 2, 6, 24 and 48 hpi (Malone *et al*. 2018). RNA-seq data from the infected bAM indicated that host genes were differentially expressed between the cells infected with each MTBC strain, suggesting that different immune response pathways may be employed by the bovine host in response to *M. bovis* and *M. tuberculosis*. In this regard, it is important to note that cattle experimentally infected with *M. tuberculosis* displayed minimal pathology, even though diagnostic assays indicated a successful infection (Whelan *et al*. 2010; Villarreal-Ramos *et al*. 2018).

In the current study, we extend our macrophage infection approach to encompass comparative integrative analyses for both pathogens (*M. bovis* and *M. tuberculosis*) and their corresponding mammalian hosts (*B. taurus* and *H. sapiens*). To do this we used four different experimental infection groups, each with parallel non-infected control cells: 1) bAM infected with *M bovis* (bAM-MB); 2) bAM infected with *M. tuberculosis* (bAM-MT); 3) hAM infected with *M. tuberculosis* (hAM-MT); and 4) human monocyte derived macrophages (hMDM) infected with *M. tuberculosis* (hMDM-MT). Transcriptomics data from these experiments were analysed, compared, and interpreted using three separate computational pipelines: 1) differentially expressed (DE) genes (DEG)—the standard approach to catalogue quantitative changes in gene expression levels between experimental groups; 2) differential gene expression interaction networks (DEN), which uses validated molecular interactions extracted from the scientific literature and other sources in combination with DE gene expression values to detect and identify functional gene subnetworks (modules); and 3) combined pathway analysis (CPA), where DE genes are subject to pathway enrichment across six different biological pathway resources. The outputs from these analyses were then integrated with two separate high-resolution bovine and human GWAS data sets with the aim of uncovering novel QTLs and obtaining new insights into the genomic architecture of the TB infection response in cattle and humans.

## 2. Materials and methods

### 2.1. Ethics statement

No animal or human procedures were performed for this computational genomics study, which used published and publicly available RNA-seq transcriptomics data.

### 2.2. General computational methods

All data-intensive computational procedures were performed on a 36-core/72-thread compute server (2× Intel^®^ Xeon^®^ CPU E5-2697 v4 processors, 2.30 GHz with 18 cores each), with 512 GB of RAM, 96 TB SAS storage (12 × 8 TB at 7200 rpm), 480 GB SSD storage, and with Ubuntu Linux OS (version 18.04 LTS).The complete computational and bioinformatics workflow is available with additional information as a public GitHub repository (<u>github.com/ThomasHall1688/Bovine_multi-</u> <u>omic_integration</u>). The individual components of the experimental and computational workflows are described below and these methodologies and procedures used are modified from those detailed previously by us (Hall *et al*. 2021).

### 2.3. Bovine genomic data acquisition

Genome-wide RNA-seq transcriptomics data we previously generated from a 48-h bAM time course challenge experiment using the sequenced *M. bovis* AF2122/97 and *M. tuberculosis* H37Rv strains was used (GEO accession: GSE62506). The complete laboratory methods used to isolate, culture and infect bAM with *M. bovis* AF2122/9 and *M. tuberculosis* H37Rv and generate strand-specific RNA-seq libraries using RNA harvested from these cells are described in detail elsewhere (Magee *et al*. 2014; Nalpas *et al*. 2015; Malone *et al*. 2018). Briefly, these RNA-seq data were generated using bAM obtained by lung lavage of ten unrelated age-matched 7–12-week-old male Holstein-Friesian calves. These bAM were infected *in vitro* with 1) *M. bovis* AF2122/97, 2) *M. tuberculosis* H37Rv, or 3) incubated with media only. Following total RNA extraction from the two MTBC strain infected groups and the control non-infected bAM, strand-specific RNA-seq libraries were prepared. These comprised *M. bovis-*, *M. tuberculosis-* and non-infected samples from each post-infection time point (2, 6, 24 and 48 hpi across 10 animals (with the exception of one animal that did not yield sufficient alveolar macrophages for *in vitro* infection at 48 hpi). Raw sequence data for RNA-seq analysis was generated as paired-end 2 × 90 nucleotide reads using an Illumina^®^ HiSeq^™^ 2000 apparatus. For the present study, the RNA-seq libraries derived from the 24 hpi timepoint for the *M. bovis*-infected bAM, *M. tuberculosis*-infected bAM, and control non-infected bAM were selected for transcriptomics analysis and downstream comparative data mining.

Bovine GWAS data sets for the present study were obtained from intra-breed imputed WGS-based GWAS analyses that used estimated breeding values (EBVs) derived from a bTB resistance phenotype, which were generated for 1,502 Holstein-Friesian sires (Ring *et al*. 2019). The bTB phenotype, the WGS-based imputed SNP data, and the quantitative genetics methods are described in detail elsewhere (Ring *et al*. 2019); however, the following provides a brief summary. The bTB resistance phenotype was defined for every animal present during each herd-level bTB breakdown when a bTB reactor or an abattoir case was identified. Cattle that yielded a positive single intradermal comparative tuberculin test (SICTT), post-mortem lymph node lesion, or laboratory culture result/s were coded as “1” (bTB = 1) and all other cattle present in the herd during the bTB-breakdown were coded as “0” (bTB = 0). After phenotype data edits, bTB resistance EBVs were generated for 781,270 cattle and these were used to produce individual sire EBVs for each of three breed groups (Charolais, Limousin and Holstein-Friesian). For the present study we used data from the Holstein-Friesian breed group. After SNP filtering using thresholds for minor allele frequency (MAF < 0.002) and deviation from Hardy-Weinberg equilibrium (HWE; *P* < 1 × 10^-6^), there were 15,017,692 autosomal SNPs for the Holstein-Friesian sire analysis. A single-SNP regression analysis was then performed using sire EBVs for bTB resistance and the nominal GWAS *P* values were used for downstream integrative genomics analyses.

### 2.4. Human genomic data acquisition

RNA-seq transcriptomics data from a 72-h hAM and hMDM time course challenge experiment using the sequenced *M. tuberculosis* H37Rv strain was used for the human transcriptomics component of this study (GEO accession: GSE114371). The complete laboratory methods used to isolate, culture, and infect hAM and hMDM with *M. tuberculosis* H37Rv and generate strand-specific AmpliSeq^™^ RNA-seq libraries using RNA harvested from these cells are described in detail elsewhere (Papp *et al*. 2018). Briefly, these RNA-seq data were generated using samples obtained from healthy human donors that tested negative for the tuberculin skin test (TST), under an approved IRB protocol at the Ohio State University Wexner Medical Centre (Papp *et al*. 2018). Human AM and PBMC (used for culturing hMDM) were obtained from different donors and macrophages (hAM and hMDM) were infected with *M. tuberculosis* H37Rv using a multiplicity of infection (MOI) of 2:1. Infected and control non-infected hAM and hMDM were harvested in Trizol reagent after 2, 24 and 72 hpi and RNA was extracted from each experimental group and Ion Torrent sequencing libraries were prepared according to the AmpliSeq^™^ Library prep kit protocol (Li *et al*. 2015).

The human GWAS data set for resistance to infection by *M. tuberculosis*, which was used for the human integrative genomics work described in this study was obtained from the UK Biobank *GeneATLAS* (http://geneatlas.roslin.ed.ac.uk). Detailed information about this public atlas of genetic associations for 118 non-binary and 660 binary traits catalogued in 452,264 UK Biobank participants of European ancestry has been published by Canela-Xandri *et al*. (Canela-Xandri *et al*. 2018) and the UK Biobank deep phenotyping and genotyping project is described by Bycroft *et al*. (Bycroft *et al*. 2018). The UK Biobank resource contains genotype data for 488,377 participants that was generated using custom high-density genome-wide SNP arrays containing more than 800,000 SNPs. These SNP data were then imputed up to more than 30 million genome-wide SNPs for the GeneATLAS GWAS resource (Canela-Xandri *et al*. 2018). The hTB GWAS data set used for the present study was generated from 2,219 hTB cases and 450,045 disease-free controls. Autosomal SNP filtering criteria consisted of the following: a SNP call rate threshold > 0.98; HWE deviation threshold (*P* < 1 × 10^-50^ on a subset of 344,057 unrelated White British individuals); MAF < 0.001; and an imputation score > 0.9 (Canela-Xandri *et al*. 2018). There were 9,113,113 SNPs remaining after these filtering steps. In common with the other traits, the binary hTB resistance trait GWAS data set was generated using a linear mixed model (LMM) and a genomic relationship matrix (GRM) using the DISSECT software tool (Canela-Xandri *et al*. 2015). **Supplementary Fig. 1** shows a Q-Q plot obtained for the expected and observed SNP *P* values for the hTB resistance trait associations.

### 2.5. Differential gene expression analysis of bovine and human RNA-seq data

Differential gene expression analysis (experimental contrast: infected versus control) of the bovine RNA-seq data, which was aligned to the bovine genome ARS-UCD1.2 (Rosen *et al*. 2020), was performed using the DESeq2 package (version 1.24.0) (Love *et al*. 2014) with a longitudinal time series design that accounted for time (hours post-infection, hpi) and treatment (control and infected). Multiple testing correction was performed using the Benjamini-Hochberg false discovery rate (FDR) method (Benjamini & Hochberg 1995). Consequently, an individual gene was considered to be differentially expressed (DE) if it exhibited an FDR-adjusted *P*-value less than 0.05 (*P*_adj._ < 0.05) and an absolute log_2_ fold-change greater than one (|log_2_FC| > 1).

The human differential gene expression analysis (experimental contrast: infected versus control) is fully described by Papp and colleagues (Papp *et al*. 2018). Briefly, human AmpliSeq^™^ RNA-seq data was analysed using the Ion Torrent Mapping Alignment Program (TMAP) (Pietrzak *et al*. 2016). Differential expression analysis was performed with the R package edgeR (Robinson *et al*. 2010). For the purposes of the work described in this study, DE genes were selected based on FDR *P*_adj._ < 0.05 and |log_2_FC| > 1.

### 2.6. Functional gene module identification using differential gene interaction networks

The GeneCards^®^ (www.genecards.org; version 4.12) gene compendium and knowledge database is a webtool that integrates multiple sources of biological information on all annotated and predicted human genes (Stelzer *et al*. 2016). This database was used to identify a set of genes that are functionally associated with the host response to diseases caused by infection with mycobacteria. The search query used was `tuberculosis` OR `mycobacterium` OR `mycobacteria` OR `mycobacterial` and genes were ranked by a GeneCards^®^ statistic—the *Relevance Score*—based on the *Elasticsearch* algorithm (Gormley & Tong 2015), which determines the strength of the relationships between genes and keyword terms. Gene IDs were converted to human Ensembl gene IDs (Martin *et al*. 2023) and retained for downstream analysis using the InnateDB knowledgebase and analysis platform for systems level analysis of the innate immune response (www.innatedb.com; version 5.4) (Breuer *et al*. 2013).

A gene interaction network (GIN) was generated with the gene list output from GeneCards^®^ using InnateDB with default settings and this network was visualised using Cytoscape (Shannon *et al*. 2003). The jActivesModules Cytoscape plugin (version 3.12.1) (Ideker *et al*. 2002) was then used to superimpose the four bovine and human RNA-seq DE genes data sets and detect—through a greedy search algorithm—differentially active subnetworks (modules) of genes. This process generated four sets of locally coherent clusters that contain both DE genes and genes that are not DE but are members of the functional gene modules. These modules were identified using: the log_2_FC and *P*_adj._ values of each DE gene; the overall connectivity of those genes with their immediate module co-members; and the comparison of that connectivity with a background comprised of randomly drawn networks using the same genes, but independent of the base network. Gene module identification revealed a set of modules for each of the four sets of DE genes (bAM infected with *M. bovis* – bAM-MB, bAM infected with *M. tuberculosis* – bAM-MT, hAM infected with *M. tuberculosis* – hAM-MT, and hMDM infected with *M. tuberculosis* – hMDM-MT). Genes embedded in active modules that were detected as statistically significant for the four experimental contrasts were combined and annotated for downstream GWAS integration.

### 2.7. Combined Pathway Analyses

For the combined pathway analysis (CPA), the four DE gene data sets were uploaded to the InnateDB pathway analysis webtool (Breuer *et al*. 2013) and bovine genes were converted to their human orthologs. Selection of genes for CPA was performed using *P*_adj._ < 0.05 and |log_2_FC| > 1, with all remaining genes acting as a background distribution by which to compare. The InnateDB overrepresentation analysis (ORA) tool was used, which performs a meta pathway analysis across multiple databases. The four different DE gene sets were queried against the following six pathway resource databases: 1) *Kyoto Encyclopaedia of Genes and Genomes* (KEGG – www.genome.jp/kegg) (Kanehisa *et al*. 2023); 2) *Integrating Network Objects with Hierarchies Pathway Database* (archived INOH – <u>dbarchive.biosciencedbc.jp/en/inoh/desc.html</u>) (Yamamoto *et al*. 2011); 3) *NCI-Nature Pathway Interaction Database* (archived NCI-PID – www.ndexbio.org) (Pillich *et al*. 2023); 4) *Reactome* (REACTOME – <u>reactome.org</u>) (Gillespie *et al*. 2022); 5) *Biocarta Pathways* (archived BIOCARTA – amp.pharm.mssm.edu/Harmonizome/dataset/Biocarta+Pathways) (Nishimura 2001); and 5) *NetPath* (NETPATH – www.netpath.org) (Kandasamy *et al*. 2010). Following this, the top five pathways from each experimental contrast were identified using InnateDB pathway overrepresentation *P*_adj._ values (B-H FDR) and curated for downstream analysis. For each of the five top pathways for the four experimental contrasts, all the genes within the pathway, regardless of whether differentially expressed, were extracted, tabulated and annotated for downstream GWAS integration.

### 2.8. Ingenuity^®^ Pathway Analysis

Ingenuity^®^ Pathway Analysis—IPA^®^ (version 1.1, summer 2020 release; Qiagen, Redwood City, CA, USA) was used to perform a statistical enrichment analysis of DE gene sets for each experimental group (Krämer *et al*. 2014). This enabled identification of canonical pathways and functional processes of biological importance in these groups. Following best practice, the background gene set for pathway and functional process enrichment testing was the set of detectable genes for each experimental group (Timmons *et al*. 2015). To produce gene sets for the IPA Core Analysis within the recommended range for the number of input entities (Krämer *et al*. 2014; Qiagen ^2^_02_^3^) and to include DE genes with small fold-change values, gene sets were filtered using only *P*_adj._ thresholds of 0.05 for the four experimental groups. In addition, the target species selected was *Homo sapiens* and the cell type used was *Macrophage* (including *Microglia* OR *Bone marrow-derived macrophages* OR *Monocyte-derived macrophages* OR *Other macrophages* OR *Peritoneal macrophages* OR *Macrophages not otherwise specified*) with the *Experimentally Observed* and *High Predicted* confidence settings. For each of the ten top pathways for the four experimental contrasts, all the genes within the pathway, regardless of whether differentially expressed, were extracted, tabulated, and annotated for downstream GWAS integration.

### 2.9. Integration with bovine and human tuberculosis GWAS data sets

To facilitate integration of GWAS data with gene sets generated from functional genomics data analyses, the *gwinteR* software package was used (<u>github.com/ThomasHall1688/gwinteR</u>) (Hall *et al*. 2021). The *gwinteR* tool can be used to test the hypothesis that a specific set of genes is enriched for signal in a GWAS data set relative to the genomic background. For example, such a gene set could be an output from an active gene module network analysis of transcriptomics data from a tissue relevant to the GWAS phenotype. For the present study, *gwinteR* was used as follows: 1) a set of SNPs (the target SNP set) was collated across all genes in a specific gene set at increasing genomic intervals upstream and downstream from each gene inclusive of the coding sequence (e.g., ±0 kb [intragenic only], ±10 kb, ±20 kb, ±30 kb… …±100 kb); 2) for each genomic interval, a null distribution of 2,500 SNP sets, each of which contains the same number of total combined SNPs as the target SNP set, was generated by resampling with replacement from the search space of the total population of SNPs in the GWAS data set; 3) the nominal (uncorrected) GWAS *P*-values for the target SNP set and the null distribution SNP sets were converted to local FDR-adjusted *P*-values (*P*_adj._) using the fdrtool R package (current version 1.2.15) (Strimmer 2008); 4) a permuted *P*-value (*P*_perm._) to the test the primary hypothesis for each observed genomic interval target SNP set was generated based on the proportion of permuted random SNP sets where the same or a larger number of SNPs exhibiting significant *q*-values (e.g. *q* < 0.05 or *q* < 0.10) are observed; 5) *gwinteR* generated data to plot *P*_perm._ results by genomic interval class and obtain a graphical representation of the GWAS signal surrounding genes within the target gene set; 6) a summary output file of all SNPs in the observed target SNP set with genomic locations and *q*-values was generated for subsequent investigation.

For the integrative analyses of GWAS data sets with functional genomics outputs from infected bAM (bAM-MB and bAM-MT) and infected hAM and hMDM (hAM-MT and hMDM-MT), four different subsets of genes for each experimental contrast were used: 1) basic DE gene sets that were filtered to ensure manageable computational loads using stringent expression threshold criteria of |log_2_FC| > 2 and *P*_adj._ < 0.01 for the bAM-MB, bAM-MT and hMDM-MT contrasts and less stringent expression criteria of |log_2_FC| > 1 and *P*_adj._ < 0.05 for the hAM-MT contrast; 2) for the four contrasts individually (bAM-MB, bAM-MT, hAM-MT, and hMDM-MT), the genes embedded in active modules identified from the GIN using jActiveModules; 3) for the four contrasts individually, all genes, regardless of expression, that are members of the top five overrepresented pathways across the KEGG, INOH, NCI-PID, REACTOME, BIOCARTA and NETPATH databases; and 4) for the four contrasts individually, all genes, regardless of expression, that are members of the top 10 enriched pathways obtained using the IPA analyses.

## 3. Results

### 3.1. Differential gene expression of *M. bovis-* and *M. tuberculosis*-infected bovine alveolar macrophages

Quality filtering of RNA-seq read pairs yielded a mean of 22,347,042 ± 2,433,115 reads per individual library (*n* = 29 libraries). A mean of 19,290,873 ± 2,166,803 read pairs (86.31%) were uniquely mapped to locations in the ARS-UCD1.2 bovine genome assembly. Detailed filtering, mapping, and read count statistics are provided in **Supplementary Information File 1** (Worksheets 1 and 2) and multivariate PCA analysis of the individual animal sample expression data using DESeq2 revealed separation of the control non-infected AM versus the bAM-MB and the bAM-MT at the 24 hpi time point (**Supplementary Fig. 2**).

Using default criteria for differential expression (*P*_adj._ < 0.05) and considering the bAM-MB and bAM-MT relative to the control non-infected AM, 3,591 DE genes were detected at 24 hpi in the bAM-MB contrast (1,879 with increased expression and 1,712 with decreased expression); and 1,816 DE genes were detected at 24 hpi in the bAM-MT contrast (1,039 increased and 777 decreased). **Fig. 1** shows volcano plots of DE genes for the bAM-MB and bAM-MT comparisons (see also **Supplementary Information File 1** – Worksheets 3 and 4).

**Fig. 1:**
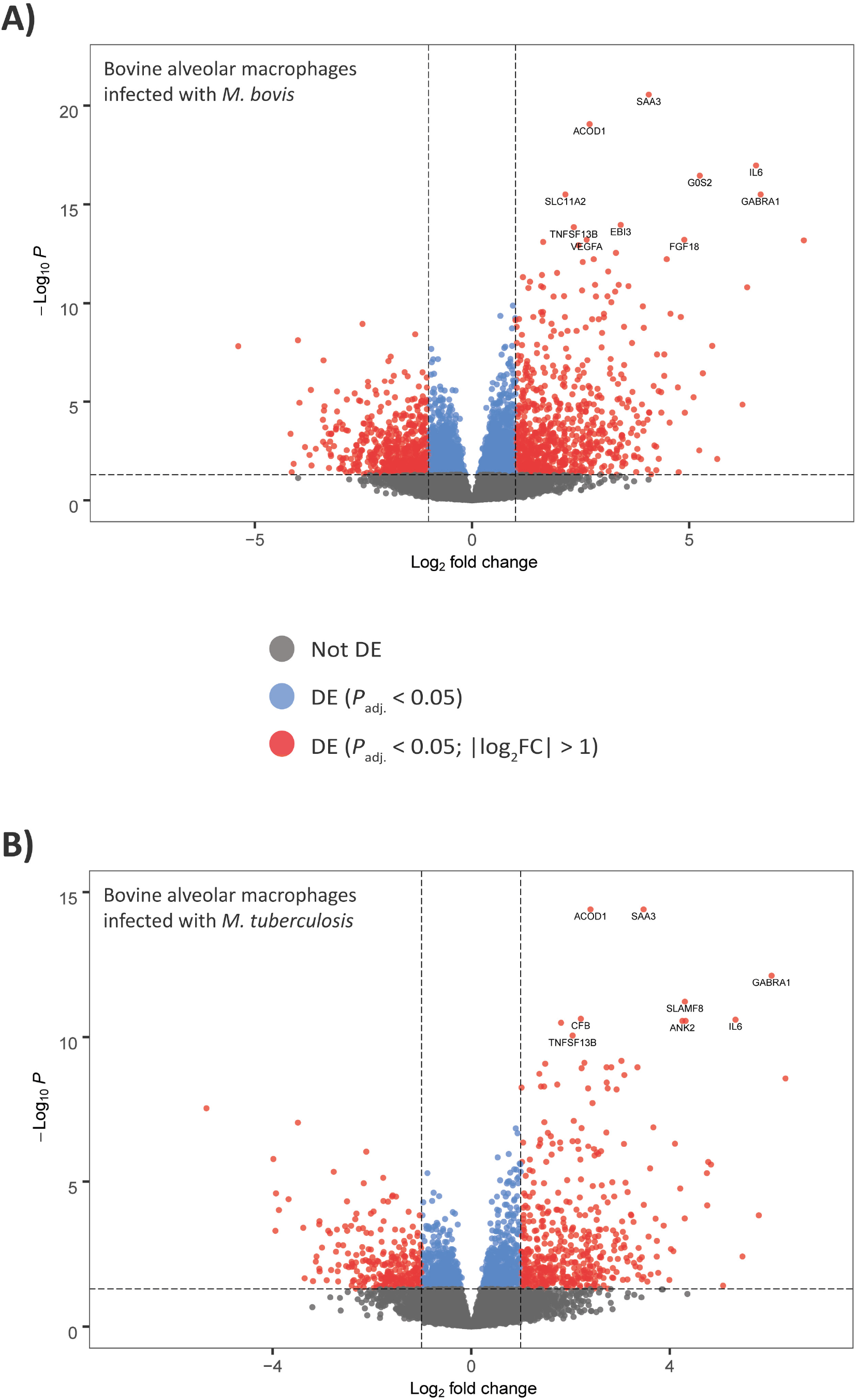
Volcano plots showing differentially expressed genes for the two bAM experimental contrasts at 24 hpi. **A.** Bovine alveolar macrophages (bAM) infected with *M. bovis* versus control non-infected bAM. **B.** bAM infected with *M. bovis* versus control non-infected bAM. All genes with *P*_adj._ < 0.05 and |log_2_FC| > 1 are shown in red, with the top ten genes by *P*_adj._ value labelled for each group.

To ensure manageable computational loads, the input lists of bAM-MB and bAM-MT DE gene sets for GWAS integration with the *gwinteR* tool (DEG-bAM-MB) and DEG-bAM-MT) were filtered with |log_2_FC| > 2, and *P*_adj._ < 0.01 and *P*_adj._ < 0.05 for the bAM-MB and bAM-MT contrasts, respectively. The less stringent *P*_adj._ threshold used for the DEG-bAM-MT input gene set was a consequence of losing one sample after QC filtering (*n* = 9). With these criteria, there were 378 and 284 genes for the DEG-bAM-MB and DEG-bAM-MT input sets, respectively. The two DEG bAM *gwinteR* input gene sets are fully detailed in **Supplementary Information File 1** (Worksheets 5 and 6). It is also important to note that 243 genes overlapped between the DEG-bAM-MB and DEG-bAM-MT gene sets.

### 3.2. Differential gene expression of *M. tuberculosis*-infected human alveolar macrophages and human monocyte-derived macrophages

A full description of the methodology used to detect DE genes in *M. tuberculosis*-infected hAM and hMDM at 24 hpi is available in the source publication (Papp *et al*. 2018). Briefly, a mean of 8,557,929 ± 2,22,249 read pairs (93.95%) from 12 libraries (3 hAM control/3 hAM infected, 3 hMDM control/3 hMDM infected) were uniquely mapped to locations in the GRCh38 human genome assembly. Detailed mapping statistics and read count information on the DE genes are provided in **Supplementary Information File 2** (Worksheets 1 and 2).

Using default criteria for differential expression (*P*_adj._ < 0.05) and considering the *M. tuberculosis*-infected hAM (hAM-MT) and hMDM (hMDM-MT) relative to the control non-infected hAM and hMDM, 899 DE genes were detected at 24 hpi for the hAM-MT contrast (567 increased and 332 decreased) and 1,545 DE genes were detected at 24 hpi for the hMDM-MT contrast (796 increased and 749 decreased). **Fig. 2** shows volcano plots of DE genes for the hAM-MT and hMDM-MT comparisons (see also **Supplementary Information File 2** – Worksheets 3 and 4).

**Fig. 2:**
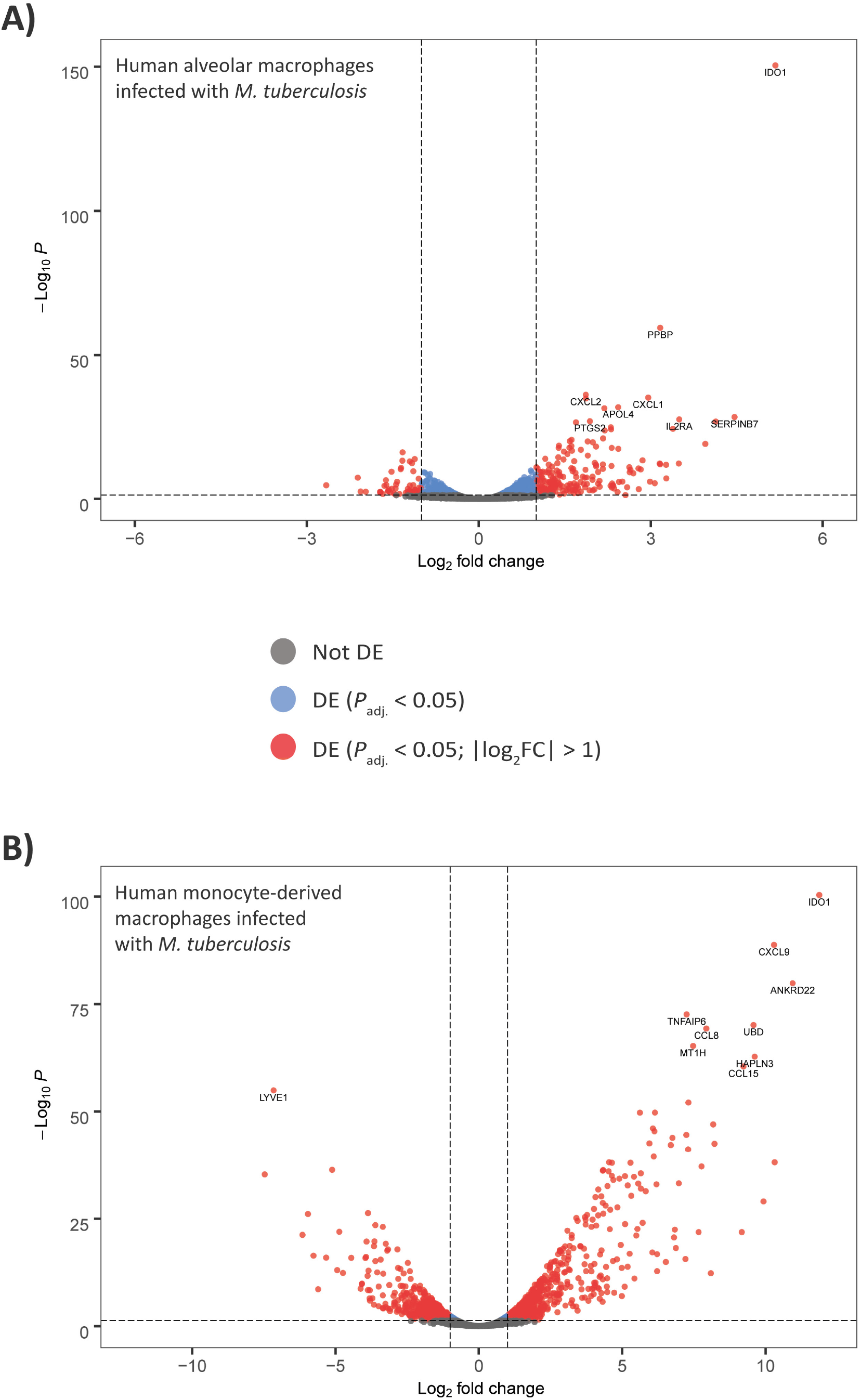
Volcano plots showing differentially expressed genes for the hAM and hMDM experimental contrasts at 24 hpi. **A.** Human alveolar macrophages (hAM) infected with *M. tuberculosis* versus control non-infected hAM. **B.** Human monocyte-derived macrophages (hMDM) infected with *M. tuberculosis* versus control non-infected hMDM. All genes with *P*_adj._ < 0.05 and |log_2_FC| > 1 are shown in red, with the top ten genes by *P*_adj._ value labelled for each group.

As with the bovine data, the input lists of hAM-MT and hMDM-MT DE gene sets that were used for GWAS integration with the *gwinteR* tool (DEG-hAM-MT) and DEG-hMDM-MT) were filtered with |log_2_FC| > 1, and *P*_adj._ < 0.05 for the hAM-MT contrast and |log_2_FC| > 2, and *P*_adj._ < 0.05 for the hMDM-MT contrast. The less stringent |log_2_FC| cut-off used for the DEG-hAM-MT input gene set was a consequence of relatively low expression fold change values observed for the hAM-MT infection experiment, which was discussed by Papp and colleagues (Papp *et al*. 2018). With these criteria, there were 277 and 415 genes for the DEG-hAM-MT and DEG-hMDM-MT input sets, respectively. The DEG-hAM-MB and DEG-hMDM-MT *gwinteR* input gene sets are fully detailed in **Supplementary Information File 2** (Worksheets 5 and 6). It is also important to note that 98 genes overlapped between the DEG-hAM-MT and DEG-hMDM-MT gene sets.

### 3.3. Common and group-specific differentially expressed genes across the four experimental contrasts

**Table 1** shows a breakdown of DE genes across the four infection groups for a range of statistical thresholds and fold-change cut-offs, including the default criteria (FDR *P*_adj._ < 0.05). The 118 up- and downregulated genes common to all experimental groups at *P*_adj._ < 0.05 are shown in **Fig. 3A** and detailed in **Supplementary Information File 2** (Worksheet 7). In addition, **Supplementary Figs. 3-6** show various between-group correlation plots for the 118 DE genes common to all experimental groups. Examples of shared upregulated genes include: *IL6*, which encodes a pro-inflammatory cytokine; *IL1B*, which also encodes an inflammatory proprotein produced by activated macrophages, *CCL20*, which encodes a chemokine that attracts dendritic cells; and *INHBA*, which encodes a member of the transforming growth factor beta superfamily. Examples of shared downregulated genes include: *PIK3IP1*, which encodes a negative regulator of *PIK3* activity, *CABLES1,* which encodes a regulator of p53/p73-induced apoptotic cell death; and *SORL1,* which encodes a transmembrane signalling receptor that is involved in control of phagocytosed mycobacteria (Vázquez *et al*. 2016). The overlap of the DE genes across the four experimental contrasts is illustrated in **Fig. 3B**.

**Fig. 3:**
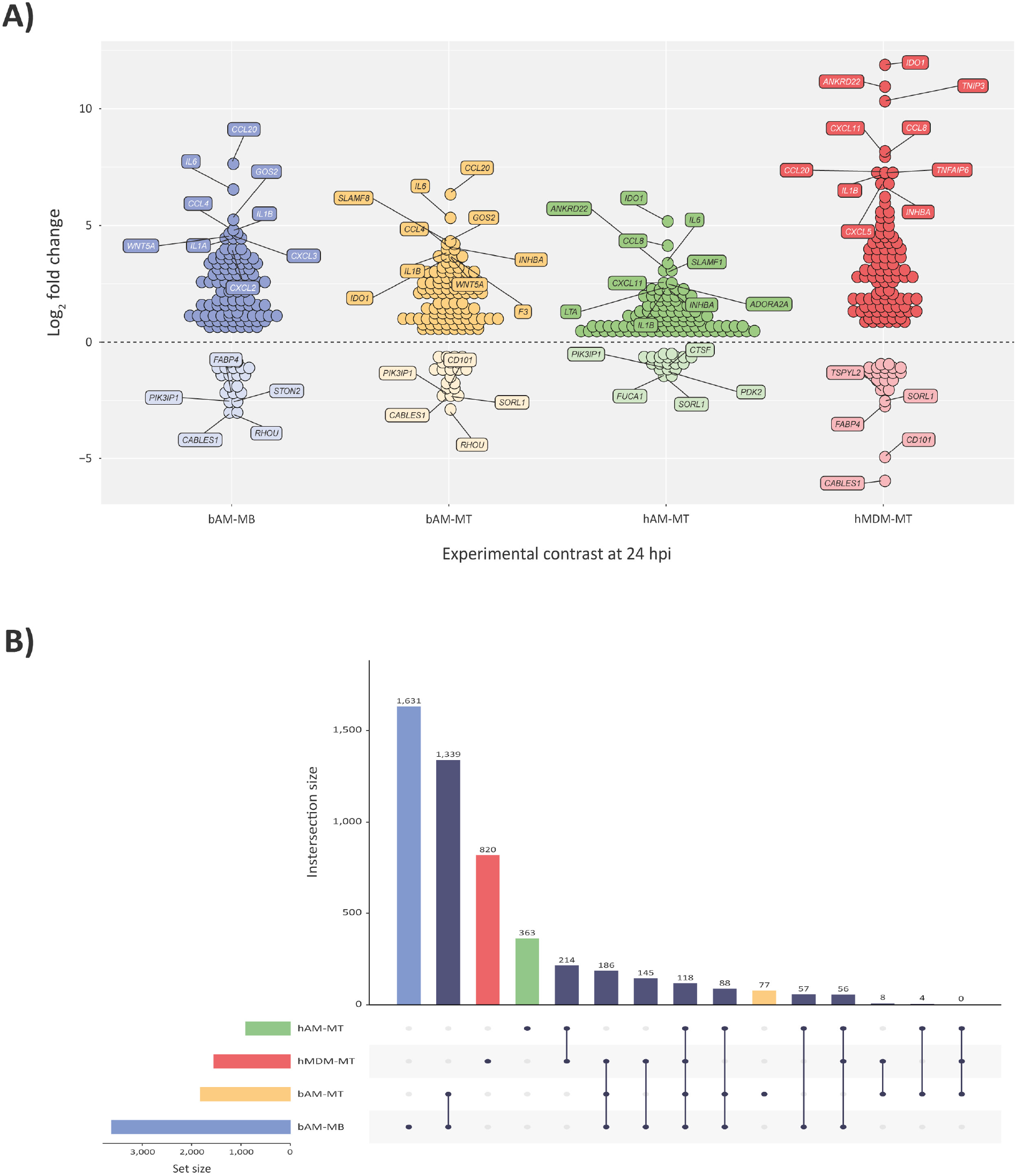
Comparison of shared differentially expressed (DE) genes across the four experimental contrasts. **A.** Plot of 118 common shared up- and downregulated genes across the four experimental contrasts. The top 10 upregulated and the top 5 downregulated genes are shown for each group (by log_2_FC). **B.** UpSetR plot showing all overlapping DE genes (*P*_adj_. < 0.05) colour-coded by experimental contrast. The UpSetR plot was generated using the UpSetR package (Conway *et al*. 2017).

**Table 1:**
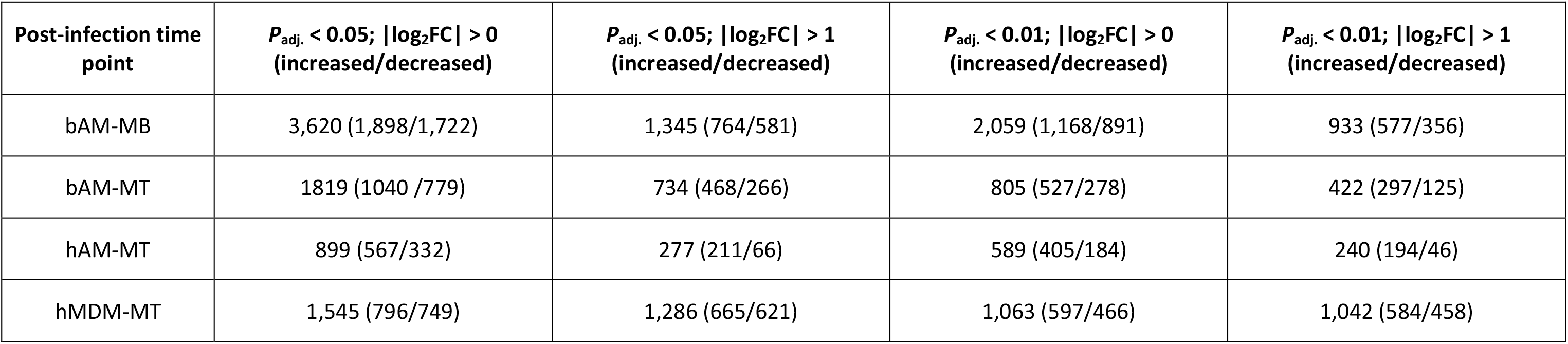
Differentially expressed genes detected in *M. bovis*-infected bovine AM (bAM-MB), *M. tuberculosis-*infected bovine AM (bAM-MT), *M. tuberculosis-*infected human AM (hAM-MT), *M. tuberculosis-*infected human MDM (hMDM-MT) relevant to controls.

### 3.4. Detection of active gene subnetworks in bovine and human infected macrophages using a tuberculosis and mycobacterial infection gene interaction network

The GeneCards^®^ search query generated a total of 2,516 gene hits using the terms `tuberculosis` OR `mycobacterium` OR `mycobacteria` OR `mycobacterial` (**Supplementary Information Files 3** and **4** – Worksheet 1). To provide a computationally manageable number of genes for an InnateDB input data set, a GeneCards^®^ relevance score (GCRS) threshold > 2.5 was used. This produced an input list of 260 functionally prioritised genes for generation of an InnateDB gene interaction network (GIN) and the top ten genes from this list ranked by GCRS were: interferon gamma receptor 1 (*IFNGR1*), toll like receptor 2 (*TLR2*), interleukin 12 receptor subunit beta 1 (*IL12RB1*), interleukin 12B (*IL12B*), solute carrier family 11 member 1 (*SLC11A1*), signal transducer and activator of transcription 1 (*STAT1*), interferon gamma receptor 2 (*IFNGR2*), cytochrome b-245 beta chain (*CYBB*), tumour necrosis factor (*TNF*), and interferon gamma (*IFNG*).

The large GIN produced by InnateDB starting with the input list of 260 functionally prioritised genes was visualised using Cytoscape and consisted of 6,951 nodes (individual genes) and 21,653 edges (gene interactions) (**Fig. 4**). Following visualisation of the large GIN in Cytoscape, the jActivesModules Cytoscape plugin was used to detect statistically significant differentially activated subnetworks (modules) within the large GIN. This consisted of superimposing the DE genes from all for experimental groups onto the larger GIN, creating differentially expressed gene interaction networks for each of the four experimental contrasts. **Supplementary Information File 3** (Worksheets 2) provides information for all gene interactions and superimposed bovine DE genes represented in **Figs. 4A** and **4B** and **Supplementary Information File 4** (Worksheets 2) provides information for all gene interactions and superimposed human DE genes represented in **Figs. 4C** and **4D**. To illustrate active module subnetwork capture, the large GINs in **Figs. 4A-D** are accompanied by an example subnetwork of genes and gene interactions. The top five subnetworks from each experimental group were retained for downstream analyses.

**Fig. 4:**
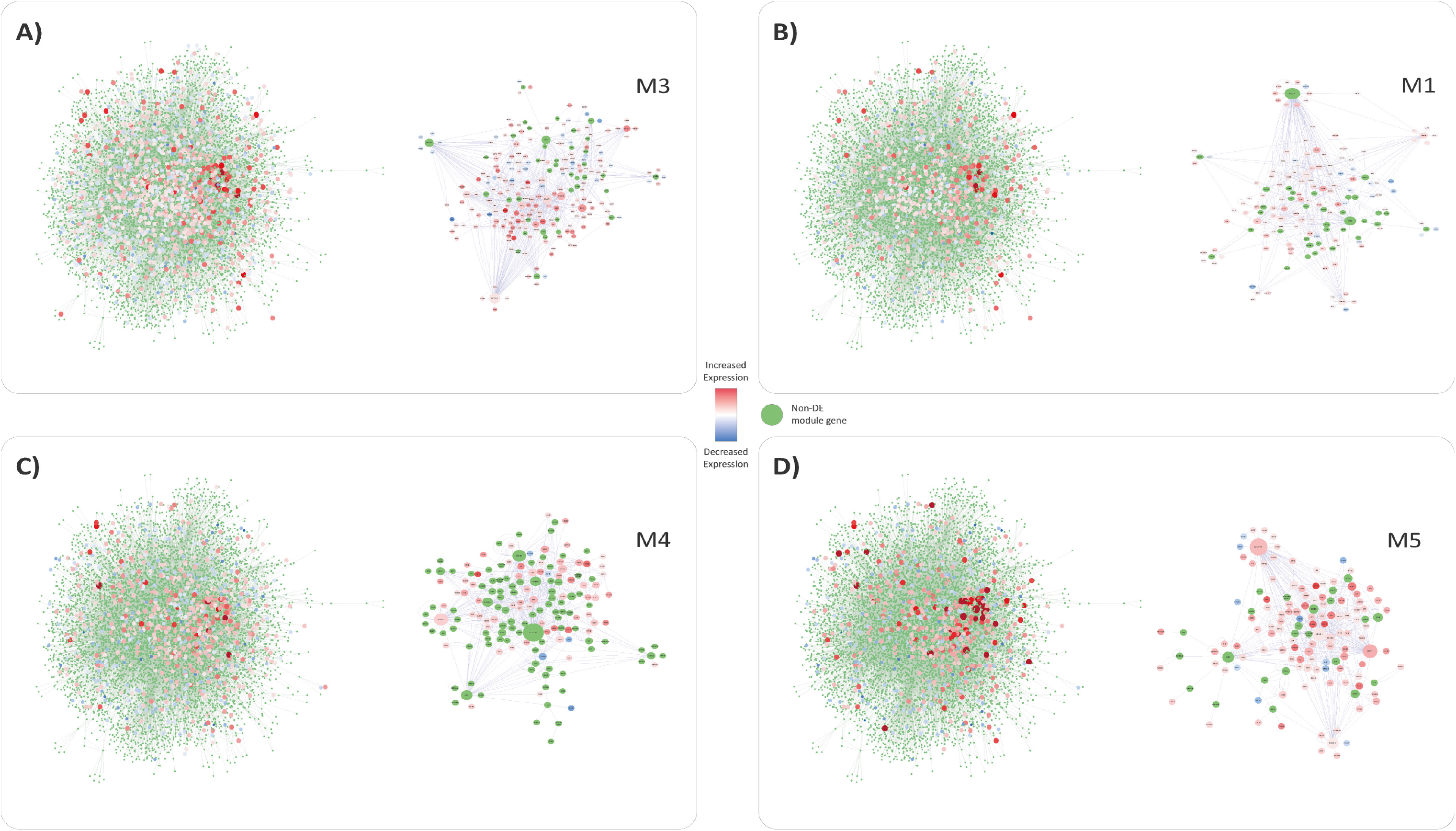
Gene interaction network analysis and functional module identification using jActiveModules and differentially expressed genes across four experimental contrasts at 24 hpi. **A.** bovine alveolar macrophages (bAM) infected with *M. bovis*. **B.** bAM infected with *M. tuberculosis*. **C.** human alveolar macrophages (hAM) infected with *M. tuberculosis*. **D.** human monocyte-derived macrophages (hMDM) infected with *M. tuberculosis*. Example active modules are shown for each contrast.

Module 1 from the bAM-MB group (M1-DEN-bAM-MB) contained 266 genes and there were 237 genes in M2-DEN-bAM-MB, 269 genes in M3-DEN-bAM-MB, 204 genes in M4-DEN-bAM-MB, and 235 genes in M5-DEN-bAM-MB (**Supplementary Information File 3** – Worksheet 3). Module 1 from the bAM-MT group (M1-DEN-bAM-MT) contained 148 genes and there were 158 genes in M2-DEN-bAM-MT, 160 genes in M3-DEN-bAM-MT, 137 genes in M4-DEN-bAM-MT, and 52 genes in M5-DEN-bAM-MT (**Supplementary Information File 3** – Worksheet 4). Module 1 from the hAM-MT group (M1-DEN-hAM-MT) contained 140 genes and there were 155 genes in M2-DEN-hAM-MT, 65 genes in M3-DEN-hAM-MT, 168 genes in M4-DEN-hAM-MT, and 180 genes in M5-DEN-hAM-MT (**Supplementary Information File 4** – Worksheet 3). Module 1 from the hMDM-MT group (M1-DEN-hMDM-MT) contained 139 genes and there were 137 genes in M2-DEN-hMDM-MT, 149 genes in M3-DEN-hMDM-MT, 93 genes in M4-DEN-hMDM-MT, and 182 genes in M5-DEN-hMDM-MT (**Supplementary Information File 4** – Worksheet 4).

After concatenation of each set of five modules and removal of duplicates, the input gene set derived from the bAM-MB group (DEN-bAM-MB) contained 398 genes and the DEN-bAM-MT input gene set contained 259 genes (**Supplementary Information File 3** – Worksheets 5 and 6). The DEN-hAM-MT input gene set contained 262 genes and the DEN-hMDM-MT input gene set contained 239 genes (**Supplementary Information File 4** – Worksheets 5 and 6).

### 3.5. Combined open-source pathway analysis of differentially expressed genes

The DE genes from each of the four experimental contrasts was analysed individually for enriched pathways across the KEGG, INOH, NCI-PID, REACTOME, BIOCARTA and NETPATH pathway repositories using the InnateDB pathway ORA tool. Pathways with a B-H FDR *P*_adj._ < 0.05 were considered enriched. Analysis of the DE genes generated from the bAM-MB experimental contrast identified 46 upregulated pathways and one downregulated pathway. After filtering by the *P*_adj._ values, the top five enriched pathways were *Cytokine-cytokine receptor interaction* (KEGG), *GPCR signalling* (INOH), *Jak-STAT signalling pathway* (KEGG), *RIG-I-like receptor signalling pathway* (KEGG), and *JAK STAT regulation* (INOH), which were all upregulated (see **Supplementary Information File 5** – Worksheet 1). Analysis of the DE genes generated from the bAM-MT experimental contrast identified 21 upregulated pathways and no downregulated pathways. The top five enriched pathways were *Cytokine-cytokine receptor interaction* (KEGG), *GPCR signalling* (INOH), *Jak-STAT signalling pathway* (KEGG), *HIF-1-alpha transcription factor network* (NCI-PID), and *IL23-mediated signalling events* (NCI-PID) (see **Supplementary Information File 5** – Worksheet 2).

Analysis of the DE genes generated from the hAM-MT experimental contrast identified 31 upregulated pathways and no downregulated pathways. The top five enriched pathways were *Cytokine-cytokine receptor interaction* (KEGG), *Chemokine receptors bind chemokines* (REACTOME), *Peptide ligand-binding receptors* (REACTOME), *Class A/1 Rhodopsin-like receptors* (REACTOME), and *GPCR ligand binding* (REACTOME) (see **Supplementary Information File 6** – Worksheet 1). Analysis of the DE genes generated from the hMDM-MT experimental contrast identified 111 upregulated pathways and three downregulated pathways. All three downregulated pathways were also significantly upregulated; however, they contained genes that were also significantly downregulated. All pathways with downregulated genes were involved in GPCR signalling. The top five enriched pathways were *Cytokine-cytokine receptor interaction* (KEGG), *Cytokine Signalling in Immune system* (REACTOME)*, Interferon gamma signalling* (REACTOME)*, Interferon Signalling* (REACTOME), and *Class A/1 (Rhodopsin-like receptors)* (REACTOME) (see **Supplementary Information File 6** – Worksheet 2). **Supplementary Fig. 7** shows the top 16 overrepresented biological pathways for each of the four experimental contrasts.

The top overrepresented pathway common to all four experimental contrasts was *Cytokine-cytokine receptor interaction* (KEGG) (see **Figs. 5** and **6**). This pathway was also the most significantly overrepresented pathway for all four experimental contrasts, though the DE genes enriched in this pathway differ between all four groups (**Figs. 5** and **6**). The genes from each of the top five pathways identified from the CPA for each experimental contrast, regardless of if the genes were DE or not, were combined, filtered for duplicates, and catalogued for GWAS integration. The CPA input gene set derived from the bAM-MB group (CPA-bAM-MB) contained 557 genes and the CPA-bAM-MT input gene set contained 473 genes (**Supplementary Information File 5** – Worksheets 3 and 4). The CPA-hAM-MT input gene set contained 640 genes and the CPA-hMDM-MT input gene set contained 737 genes (**Supplementary Information File 6** – Worksheets 3 and 4).

**Fig. 5:**
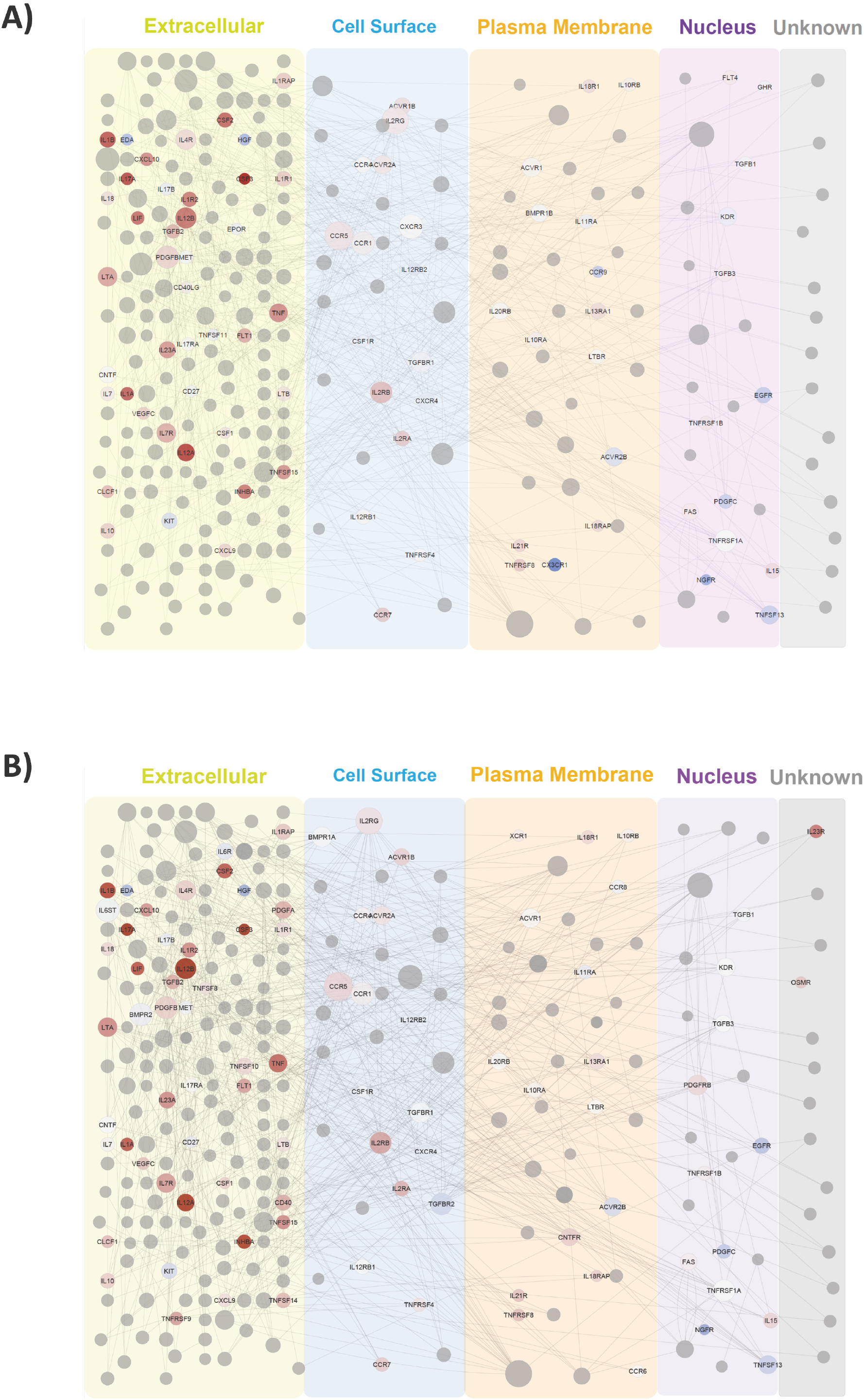
Protein network schematics and cellular locations of the top overrepresented common biological pathway - *Cytokine-cytokine receptor signalling* (bovine). **A.** *M. bovis*-infected bAM. **B.** *M. tuberculosis*-infected bAM. Red and blue nodes indicate upregulation and downregulation, respectively. Grey nodes indicate no detectable expression change. The size of the nodes corresponds to the degree of connectivity (number of interactions).

**Fig. 6:**
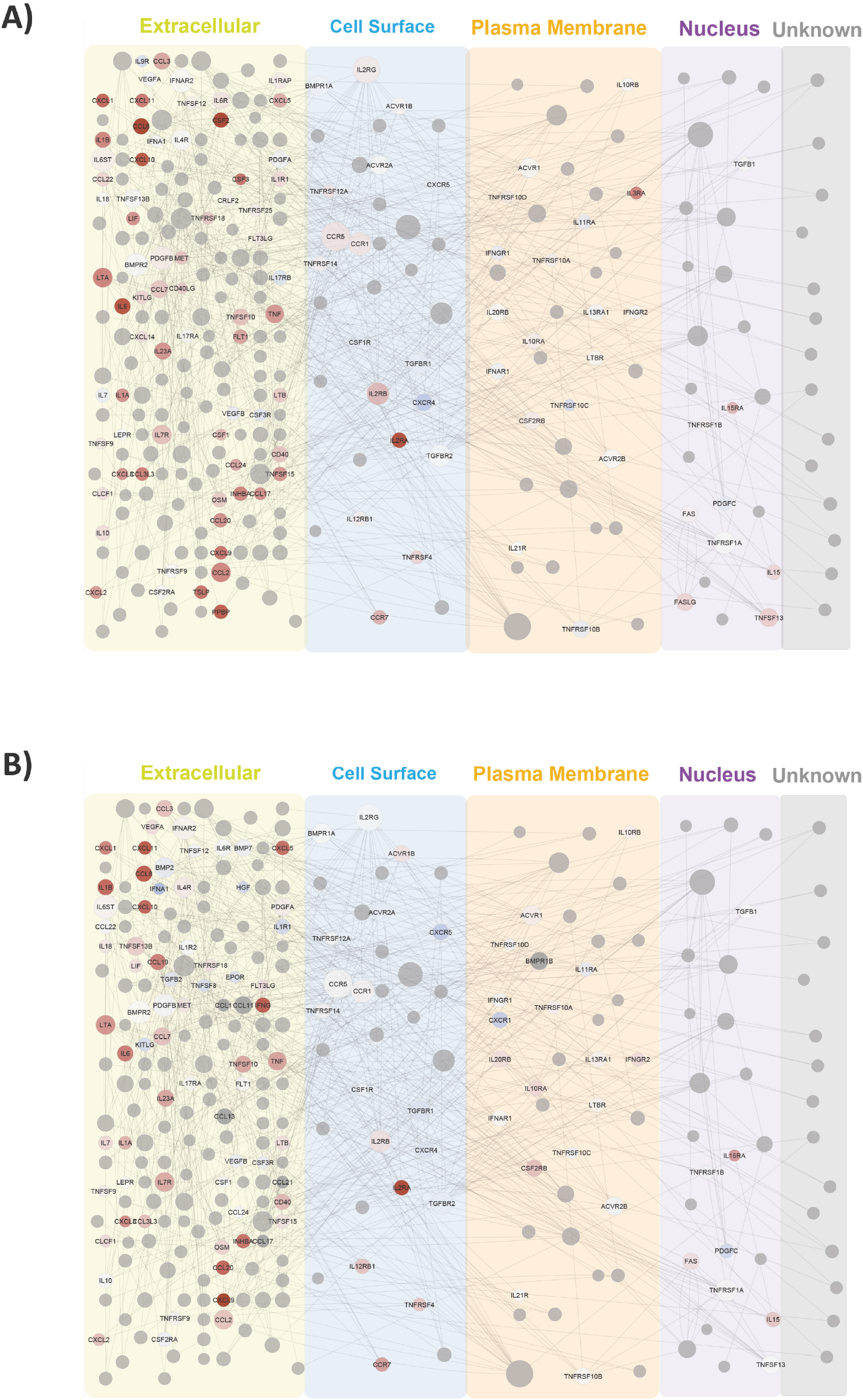
Protein network schematics and cellular locations of the top overrepresented common biological pathway – *Cytokine-cytokine receptor signalling* (human). **A.** *M. tuberculosis*-infected hAM. **B.** *M. tuberculosis*-infected hMDM. Red and blue nodes indicate upregulation and downregulation, respectively. Grey nodes indicate no detectable expression change. The size of the nodes corresponds to the degree of connectivity (number of interactions).

### 3.6. Ingenuity^®^ Pathway Analysis (IPA) of differentially expressed genes

The DE genes from each experimental contrast were analysed for enriched pathways using IPA and the IPA Knowledge Base. This resulted in 1,961 input genes (1,073 upregulated and 888 downregulated) from a background detectable set of 7,103 from bAM infected with *M. bovis* (bAM-MB), 978 input genes (571 upregulated and 407 downregulated) from a background detectable set of 6,970 genes from bAM-MT, 581 input genes (354 upregulated and 227 downregulated) from a background detectable set of 6,627 genes from hAM-MT and 941 input genes (509 upregulated and 432 downregulated) from a background detectable set of 6,825 genes from hMDM-MT.

Using the B-H method for multiple test correction in IPA (FDR *P*_adj._ < 0.05), there were 68 statistically significant enriched IPA canonical pathways from the bAM-MB group, 201 from the bAM-MT group, 61 from the hAM-MT group, and 118 from the hMDM-MT group. The genes from each of the top 10 pathways for each of the four experimental groups (**Supplementary Information File 7** and **8** – Worksheets 1 and 2), regardless of if the genes were DE or not, for each group from the IPA analysis were combined, duplicates removed, and catalogued for data integration using the bovine GWAS and human GWAS data sets (**Supplementary Information File 7** and **8** – Worksheets 3 and 4). The IPA gene set derived from bAM infected with *M. bovis* (IPA-bAM-MB) contained 386 genes and the IPA-bAM-MT input set contained 276 genes (**Supplementary Information File 7** – Worksheets 3 and 4). The IPA gene set derived from hAM infected with *M. tuberculosis* (IPA-hAM-MT) contained 207 genes and the IPA-hMDM-MT input set contained 274 gene (**Supplementary Information File 8** – Worksheets 3 and 4). **Fig. 7** summarises the top 10 IPA enriched pathways for each of the four experimental contrasts.

**Fig. 7:**
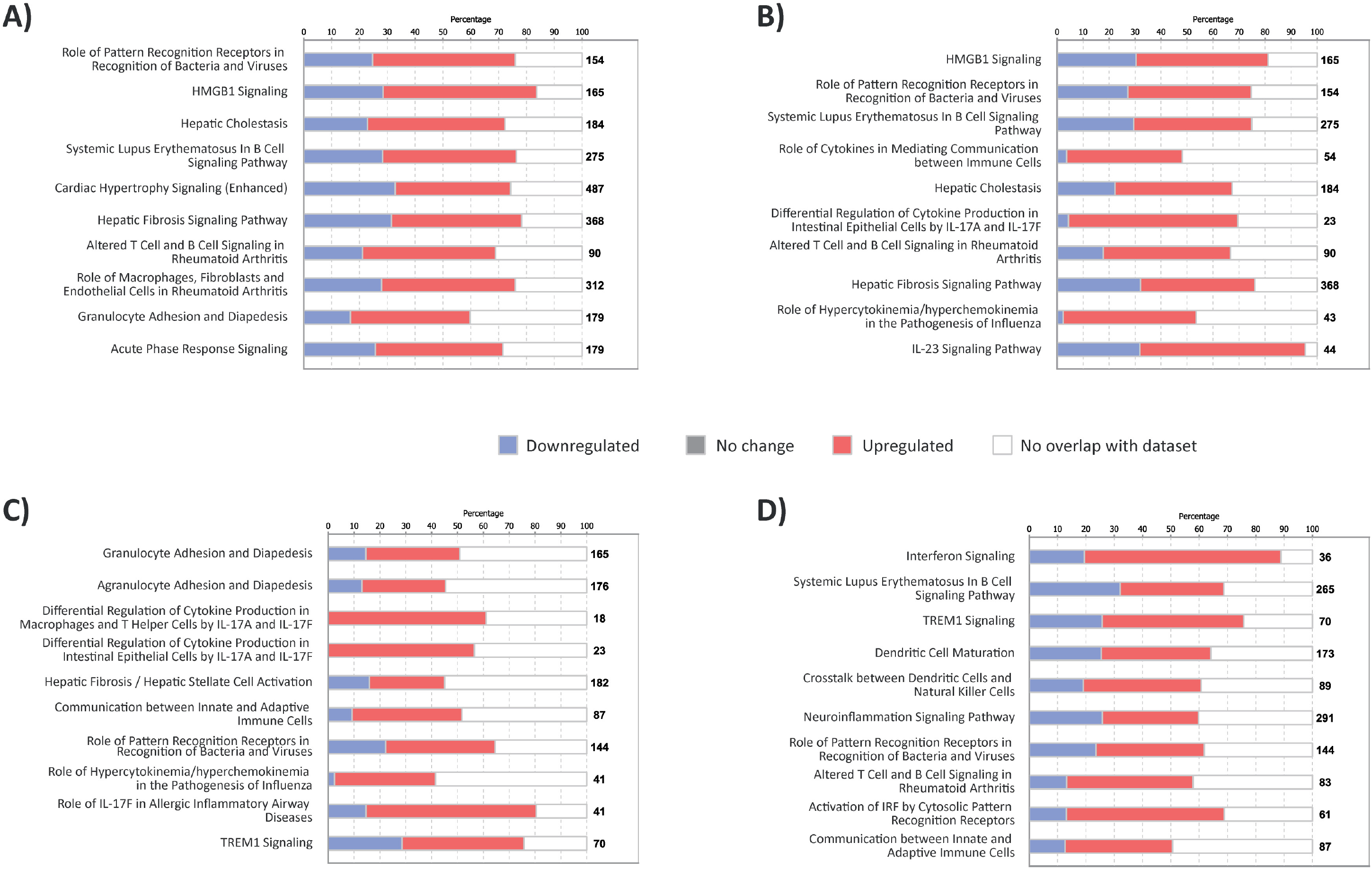
Top 10 enriched biological pathways for each experimental group from the Ingenuity^®^ Pathway Analysis (IPA). Stacked horizontal bar charts showing the top twenty canonical pathways from IPA in order of descending -log_10_ *P*_adj._: **A.** bAM-MB, **B.** bAM-MT, **C.** hAM-MT, and **D.** hMDM-MT The numbers to the right of the bars show the number of genes in the pathway and the colours of the bars indicate the percentages of these genes that are downregulated, upregulated, show no change in expression, or have no overlap with the data set.

Some of the top 10 enriched pathways (**Fig. 7**) were common to two or more experimental groups (see **Supplementary Information File 8** – Worksheet 5) and include: *Role of Pattern Recognition Receptors in Recognition of Bacteria and Viruses*; *Activation of IRF by Cytosolic Pattern Recognition Receptors; TNFR2 Signalling; Dendritic Cell Maturation; Role of Macrophages, Fibroblasts and Endothelial Cells in Rheumatoid Arthritis;* and *Hepatic Cholestasis*.

### 3.7. Integration of functional genomics outputs with GWAS data sets and identification of additional species-specific SNP-trait associations

The sixteen input gene sets generated from the four separate analyses of DE genes generated from the bAM-MB, bAM-MT, hAM-TB, and hMDM-TB infection challenge experiments are summarised in **Table 2** and further detailed in **Supplementary Information Files 1-8**. In addition to these sixteen putative functionally relevant gene sets, 100 sets of 250 genes randomly sampled from the bovine genome (BOV-RAN), and 100 sets of 250 genes randomly sampled from the human genome (HUM-RAN), were used for statistical context and comparison (data not shown). The results from the integrative analyses using the *gwinteR* tool with the DEG-bAM-MB, DEG-bAM-MT, DEN-bAM-MB, DEN-bAM-MT, CPA-bAM-MB, CPA-bAM-MT, IPA-bAM-MB, IPA-bAM-MT, and BOV-RAN input gene sets are summarised graphically in **Fig. 8** and fully detailed in **Supplementary Information File 9** (Worksheets 1-4). Likewise, the results from the integrative analyses using the equivalent human data sets are summarised graphically in **Fig. 9** and fully detailed in **Supplementary Information File 10** (Worksheets 1-4). Inspection of **Fig. 8B** and **8C** shows that, in terms of SNP enrichment (*P*_perm._ < 0.05), the integrative analyses using *gwinteR* were most effective with the DEG-bAM-MB, DEG-bAM-MT, and CPA-bAM-MT data sets. A possible explanation for the inferior performance of the other data sets is that these methods rely heavily on gene orthology, i.e., all the bovine DE genes used to generate the DEN, CPA and IPA gene sets were first converted to human gene IDs. In this regard, methods that do not rely on ortholog conversion, such as gene co-expression networks, seem to perform better. **Fig. 9B** and **9C** demonstrate that the human data integration performed better with multiple techniques; statistically significant SNP enrichment was achieved with the DEG-hAM-MT, DEG-hMDM-MT, DEN-hAM-MT, and CPA-hAM-MT data sets. **Fig. 10** provides information on the 32 genes that were proximal to SNPs significantly associated with bTB and hTB disease resilience, which were identified through integration of functional genomics outputs with the bovine and human GWAS data sets.

**Fig. 8:**
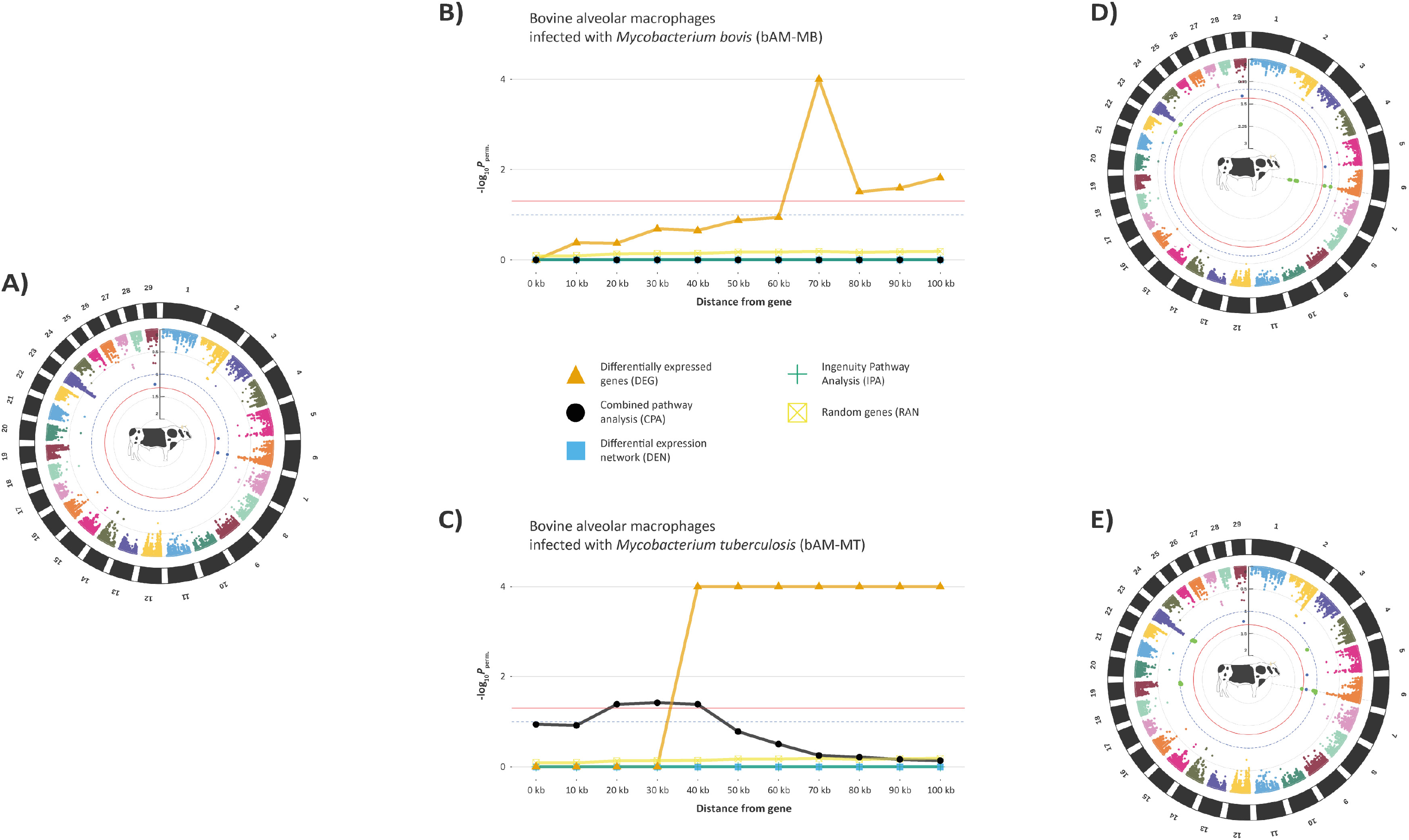
Integration of bAM functional genomics outputs and GWAS data for *M. bovis* infection resistance in Holstein-Friesian cattle. **A.** Circular Manhattan plots showing GWAS results pre-integration with blue and red highlighted data points indicating binned SNP clusters with FDR *P*_adj._ < 0.10 and < 0.05, respectively. **B.**/**C.** Line plots of permuted *P* values (-log_10_*P*_perm._) across different genomic intervals for SNPs from eight different input gene sets and random genes (**B.** bAM-MB and **C.** bAM-MT). **D.**/**E.** Circular Manhattan plots showing GWAS results post-integration with blue and red data highlighted points indicating binned SNP clusters with FDR *P*_adj._ < 0.10 and < 0.05, respectively, and SNPs enriched by *gwinteR* coloured green (**D.** bAM-MB and **E.** bAM-MT).

**Fig. 9:**
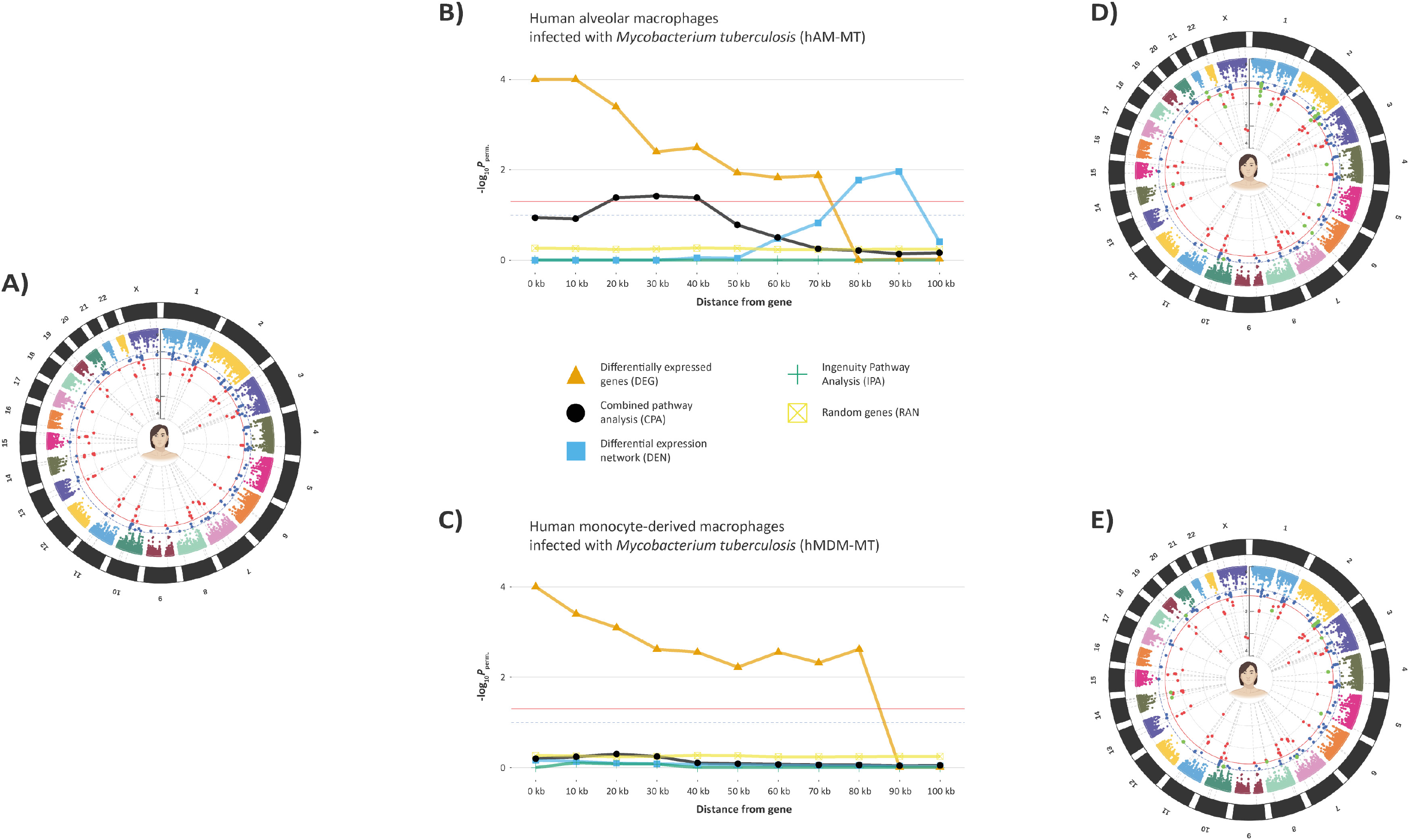
Integration of hAM and hMDM functional genomics outputs and GWAS data for *M. tuberculosis* infection resistance in humans. **A.** Circular Manhattan plots showing GWAS results pre-integration with blue and red highlighted data points indicating binned SNP clusters with FDR *P*_adj._ < 0.10 and < 0.05, respectively. **B.**/**C.** Line plots of permuted *P* values (-log_10_*P*_perm._) across different genomic intervals for SNPs from eight different input gene sets and random genes (**B.** hAM-MT and **C.** hMDM-MT). **D.**/**E.** Circular Manhattan plots showing GWAS results post-integration with blue and red data highlighted points indicating binned SNP clusters with FDR *P*_adj._ < 0.10 and < 0.05, respectively, and SNPs enriched by *gwinteR* coloured green (**D.** hAM-MT and **E.** hMDM-MT).

**Fig. 10:**
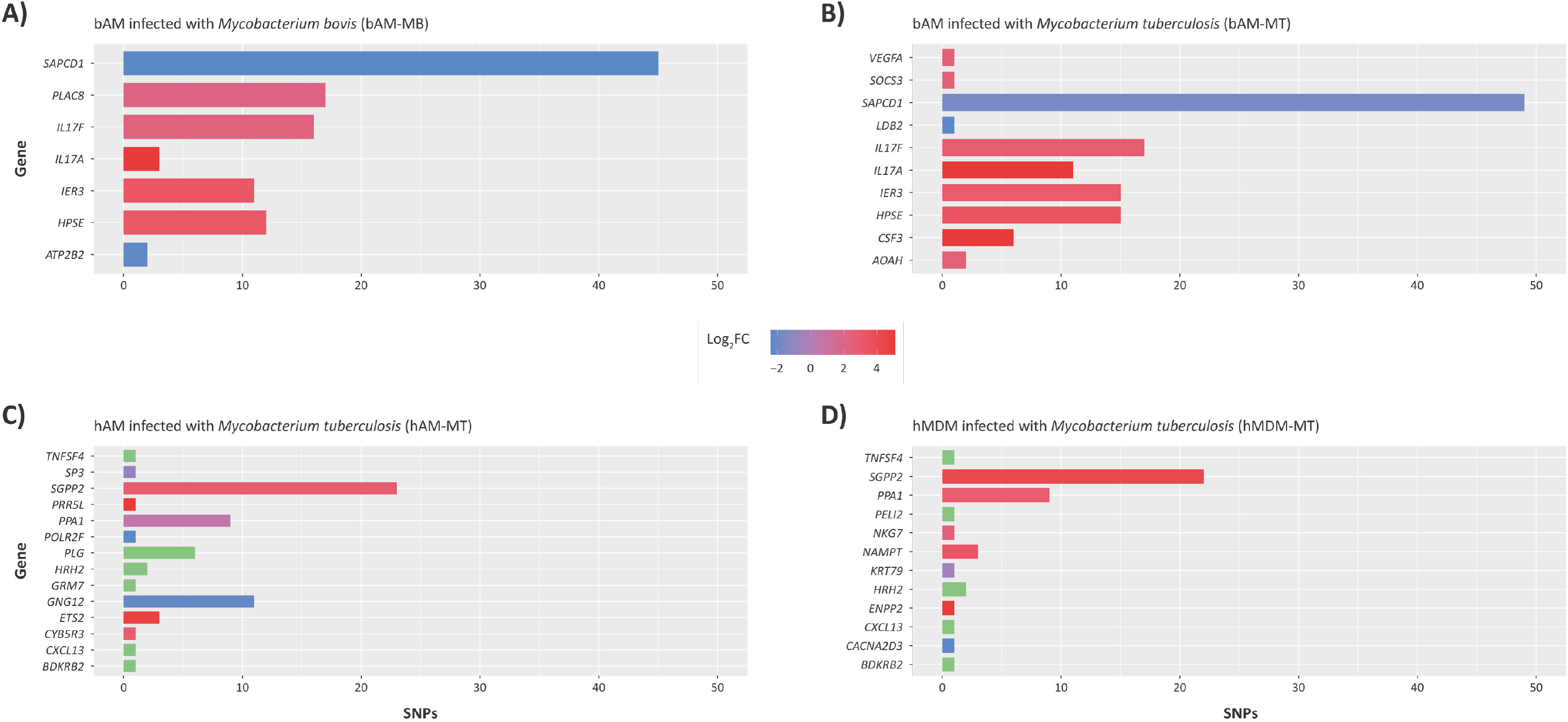
Histograms of statistically significant SNP numbers and proximal genes enriched through integration of bovine and human functional genomics outputs with bTB and hTB GWAS data sets. **A.** bAM infected with *M. bovis* (bAM-MB). **B.** bAM infected with *M. tuberculosis* (bAM-MT). **C.** hAM infected with *M. tuberculosis* (hAM-MT). **D.** hMDM infected with *M. tuberculosis* (hMDM-MT). The colour of each bar indicates whether the gene was upregulated (red), downregulated (blue), or whether there was no significant differential expression for genes identified through the subnetwork (module) or pathway analyses.

**Table 2:** The 16 different input bovine and human gene sets used for GWAS data integration.

| Infection group | Differentially expressed genes (DEG) |  | Differential expression network (DEN) |  | Combined Pathway Analysis (CPA) |  | Ingenuity Pathway Analysis (IPA) |  |
| --- | --- | --- | --- | --- | --- | --- | --- | --- |
|  | Input gene set code | No. of genes | Input gene set code | No. of genes | Input gene set code | No. of genes | Input gene set code | No. of genes |
| Bovine alveolar macrophages infected with *M. bovis* | DEG-bAM-MB | 378 | DEN-bAM-MB | 398 | CPA-bAM-MB | 557 | IPA-bAM-MB | 386 |
| Bovine alveolar macrophages infected with *M. tuberculosis* | DEG-bAM-MT | 390 | DEN-bAM-MT | 259 | CPA-bAM-MT | 473 | IPA-bAM-MT | 276 |
| Human alveolar macrophages infected with *M. tuberculosis* | DEG-hAM-MT | 277 | DEN-hAM-MT | 262 | CPA-hAM-MT | 473 | IPA-hAM-MT | 207 |
| Human monocyte-derived macrophages infected with *M. tuberculosis* | DEG-hMDM-MT | 415 | DEN-hMDM-MT | 239 | CPA-hMDM-MT | 737 | IPA-hMDM-MT | 274 |

## 4. Discussion

### 4.1. Bovine and human macrophages share common immune response genes and pathways to mycobacterial infection

The work described in this study has generated new scientific information regarding host-pathogen interaction during the initial stages of infection for *M. bovis* and *M. tuberculosis,* which are the archetypal mycobacterial pathogens of cattle and humans, respectively. Using four separate multi-omics analysis workflows, we demonstrate that there are striking similarities in the patterns of differential expression and the cellular pathways that are perturbed by these infections in bovine and human macrophages. In terms of understanding host-pathogen interaction in an evolutionary context, it is now considered likely that the MTBC complex emerged relatively recently during the Neolithic period and probably within the last 5,000 years (Bos *et al*. 2014; Kay *et al*. 2015; Sabin *et al*. 2020). Therefore, the recent shared evolutionary histories of *M. bovis* and *M. tuberculosis* may account for congruent host responses from the pathogen perspective. However, from the perspective of the mammalian host species, what is particularly remarkable—given the similarities in macrophage responses to infection—is that artiodactyls and primates last shared a common ancestor more than 80 million years ago (Liu *et al*. 2017).

As shown in **Fig. 3** and **Supplementary Information File 2** (Worksheet 7), there were 118 DE genes in common across the four different experimental infection groups (bAM-MB, bAM-MT, hAM-MT, and hMDM-MT) 24 hpi. All these genes were significantly DE, and the log_2_ fold change values were highly correlated for each pairwise comparison, with the highest correlation coefficient for these 118 genes observed between the bAM-MB and bAM-MT contrasts (*r* = 0.9894) (**Supplementary Fig. 3**). The correlation of expression differences for these genes between the hAM-MT and hMDM-MT infection groups was lower (*r* = 0.8088) (**Supplementary Fig. 4**), reflecting cell type differences in the transcriptomes of these human macrophages infected with *M. tuberculosis*. The hAM-MT/bAM-MB and hAM-MT/bAM-MT infection group comparisons for these 118 shared DE genes produced correlation coefficients of *r* = 0.7122 and *r* = 0.7170, respectively (see **Supplementary Figs. 5** and **6**); the interspecific comparison with the same MTBC pathogen (*M. tuberculosis*) being marginally more correlated.

Several groups of related genes were consistently upregulated for all four experimental infection contrasts, suggesting a core MTBC infection response gene repertoire intrinsic to macrophages from both species of mammal. Several of these genes encode different types of cytokines including interleukins (*IL1A, IL1B, IL23, IL6,* and *IL7R*); chemokines (*CCL2, CCL3, CCL4, CCL8, CCL20, CXCL2, CXCL3, CXCL5* and *CXCL11*); and members of the TNF family, such as *TNF, CD40, LTA* and *TNFAIP6*. In this regard, many of these genes were previously identified by our group as corresponding to a core innate immune response for bovine AM infected with both *M. bovis* and *M. tuberculosis* across a 48-h post-infection time course (Malone *et al*. 2018). Interestingly, the group of cytokines that did exhibit a marked species-specific pattern of gene expression were interferons. Many interferon-induced genes were differentially expressed in each experimental infection contrast group (bAM-MB: 18, bAM-MT: 6, hAM-MT: 12, and hMDM-MT: 14); however, only *IFIT2* was DE across all four groups, suggesting that for MTBC infections of AM or MDM, interferon-induced proteins tend to function in a strain- or host species-specific fashion, as opposed to interleukins and chemokines, which were consistently DE across the experimental groups regardless of macrophage cell type, strain or host species.

Other genes that were DE across the four experimental contrasts included: *STAT1*, which was consistently upregulated and is a key modulator of the immune response to mycobacterial infection and a critical transcription factor in the JAK-STAT and interferon signalling pathway (Yi *et al*. 2020); *PIK3IP1*, which was consistently downregulated and is a key negative regulator of the PIK3-AKT signalling pathway (DeFrances *et al*. 2012; Hall *et al*. 2020); and *IDO1*, which was consistently upregulated and is involved with limitation and accumulation of tryptophan in immune cells, which can induce apoptosis, limit growth of intracellular pathogens, and control immunopathology resulting from unchecked immune responses (van Baren & Van den Eynde 2015). Recent studies have indicated that the IDO1 protein may facilitate bacterial survival in macrophages infected with *M. tuberculosis*, and that downregulation of *IDO1* may lead to enhanced control of infection (Guo *et al*. 2019; Singh *et al*. 2023). It is therefore noteworthy in the present study that *IDO1* is uniformly upregulated 24 hpi across all experimental groups (bAM-MB, linear fold change of +13.3; bAM-MT, 12.3-fold; hAM-MT, +36.1-fold; and hMDM-MT, +3765.6-fold) (see Supplementary File 2 – Worksheet 7). Finally, *SLAMF1*, which was consistently upregulated and induces expression of *IL12* and *TNF* in the presence of LPS (Wang *et al*. 2004), and regulates phagosome maturation and recruitment of the PI3K complex 2; for example, with *Escherichia coli* infection of macrophages this occurs through recognition of OmpC and OmpF on the bacterial cell surface (Berger *et al*. 2010; Ma *et al*. 2012). Our group has also recently shown that the activity of *SLAMF1* is regulated at the epigenetic level, with H3K4me3 deposition at the transcriptional start site of *SLAMF1* modulating its expression in bovine AM infected with *M. bovis* (Hall *et al*. 2020).

In addition to common response genes, all four experimental infection contrasts had the same top response pathway, *Cytokine-cytokine receptor interaction* (**Figs 5** and **6**). While the DE genes in each pathway varied among the four groups, this pathway was statistically highly enriched across all the groups (**Supplementary Fig. 7**), which again highlights the evolutionary conservation and functional importance of cytokine signalling during MTBC infections of mammalian macrophages.

As part of the differential expression network analysis (DEN), a large gene interaction network (GIN) was used with log_2_ fold-change and FDR *P*_adj._ values as the key parameters to identify and extract smaller functional modules/subnetworks (see **Fig. 4** and **Supplementary Information Files 3** and **4)**. Five functional modules were extracted from the large GIN for each of the four experimental infection contrasts. Each of these subnetworks exhibits a scale-free topology (Barabasi & Albert 1999; Albert 2005); most gene nodes within the network interact with one other gene (low degree), while a small subset interacts with substantially more (high degree). These nodes tend to be direct regulators of other genes such as transcription factors, or subunits of important proteins. Comparison of genes with high degree across all 20 modules (**Supplementary Information File 4** – Worksheet 7), demonstrated that all four groups share genes of high degree, suggesting that the key regulators of gene modules that respond to intracellular mycobacterial infection are common across the two species and macrophage cell type and MTBC strain. Genes present in at least one functional module from all experimental infection contrasts included inflammation-related transcription factors such as *CEBPB, EGR1, IRF1, NFKB1, NFKBIA, STAT1, JAK2, UBC*, and *TNF*. In this regard, using a differential network approach to analyse microarray gene expression data from bovine MDM challenged with *M. bovis* and an attenuated *M. bovis* BCG vaccine strain, our group previously identified *NFKB1* and *EGR1* as key hub and bottleneck gene nodes, respectively (Killick *et al*. 2014).

Interestingly, although the *IRF1* gene was present in all experimental infection contrasts, and *IFNG, IFNG1, IFNG2, IFNG3* and *IFNG4* were present as high degree genes in some contrasts, only *IFNG* was present in more than one group and no interferon gene was present in the functional modules identified for the bAM-MT infection contrast. This observation indicates that interferon genes in these experimental contrast exhibit specific functions depending on cell type, species and MTBC strain. There was also one other gene that did not appear in subnetworks identified for the bAM-MB infection contrast, but that was present in the remaining three *M. tuberculosis*-challenged groups (bAM-MT, hAM-MT, and hMDM-MT). This was the vitamin D receptor gene (*VDR*), which mediates the immunological function of vitamin D3, an activator of macrophages (Weiss & Schaible 2015; BoseDasgupta & Pieters 2018). Vitamin D deficiency has been implicated in susceptibility to both bTB and hTB (Nelson *et al*. 2012; Hu *et al*. 2016); therefore, it is somewhat surprising that *VDR* was not a gene node in any of the modules from the bAM-MB infection contrast. Taking into consideration the inter- and intra-species robustness of the functional modules, further research is needed to improve identification and functional annotation of gene response modules extracted from knowledge-based gene interaction networks.

### 4.2. Genes enriched for bTB and hTB GWAS SNPs in both species play central roles in NF-κB signalling and the formation of the granuloma

After all of the input gene sets were integrated with the bTB and hTB GWAS data sets using *gwinteR*, a total of 32 unique genes (12 bovine and 20 human genes; see **Fig. 10**) exhibited intragenic SNPs or SNPs within 100 kb up- and downstream that were enriched for associations with infection susceptibility/resistance. There was intraspecies overlap in enriched genes, such as bovine *IL17A* and human *CXCL13*, but there was no interspecies overlap. However, it is important to note that there was an enrichment observed for genes involved with initiation, formation, and regulation of the granuloma in both species.

Granulomas are densely compact, organized aggregates of immune cells consisting of epithelioid cells, blood-derived infected and uninfected macrophages, foamy macrophages, and multinucleated giant cells (Guirado & Schlesinger 2013). The enriched bovine SNPs lay in proximity to six genes related to granuloma biology. These are *CSF3, HPSE, IER3, IL17A, IL17F* and *VEGFA*. The HPSE protein (heparanase) has a role in inflammation and cell adhesion during granuloma formation and is also expressed in peripheral granulomas (Irony-Tur-Sinai *et al*. 2003; Elad *et al*. 2013). Expression of *IER3* has been observed in chronic lung granulomas (Mehra *et al*. 2013), and it may act as an inhibitor or contributor to their formation. *VEGFA* expression in macrophages regulates granuloma formation in the non-angiogenic pathway during TB disease and recruits immune cells to the granuloma (Harding *et al*. 2019).

The *CSF3* gene (aka *GCSF*) encodes a protein with many immunological roles, such as survival, proliferation, and polarization of macrophages (Hollmén *et al*. 2016; Wen *et al*. 2019); it has also been detected in multiple macrophage infection studies using pathogenic MTBC strains (Malone *et al*. 2018; Papp *et al*. 2018; Hall *et al*. 2020). Although it has been shown to be upregulated in sarcoidosis granulomas (Casanova *et al*. 2020), the role of CSF3 in TB-induced granuloma formation has not been extensively studied; however, induction of granulomatous tissue by CSF3 has been documented in other conditions, such as granulomatous dermatitis and chronic granulomatous disease in immunocompromised patients (Sekhsaria *et al*. 1996; Ozaki *et al*. 2015). However, it is important to note that colony stimulating factor proteins have recently been shown—in a membrane fusion-independent manner—to stimulate the formation of multinucleated giant cells (MGCs), which are key components of TB-induced granulomas. For example, CSF1 in conjunction with a persistent ligand, such as TNF, can induce MGCs from bone marrow myeloid progenitors via polyploidy, typically associated with replication stress or DNA damage (Herrtwich *et al*. 2018), which supports a larger role for CSF proteins in the formation of TB granulomas. In addition, our group has recently shown that the *CSF3* locus is epigenetically regulated through H3K4me3 deposition in bovine AM infected with *M. bovis* (Hall *et al*. 2020). Consequently, based on the multi-omics results presented here that highlight *CSF3* in macrophage responses to mycobacterial infections, further research into the role of *CSF3* in bTB and hTB is warranted.

For the human functional genomics outputs and GWAS data integration results, the enriched SNPs identified were proximal to five genes associated with granuloma biology: *CACNA2D3, CXCL13, ENPP2, PLG,* and *TNFSF4*. Differential levels of CACNA2D3, a protein in the voltage-dependent calcium channel complex, had been demonstrated within granulomatous structures (Crouser *et al*. 2017) and it has been proposed to play a role in the CRR5 pathway in macrophages, which regulates chemotaxis during inflammatory responses (Shaheen *et al*. 2019). The CXCL13 chemokine recruits B cells to sites of inflammation, including the granuloma (Khader *et al*. 2009; Armas-González *et al*. 2018). Macrophages do not typically express this gene, which would explain the lack of differential expression observed in any of the four experimental infection contrasts. However, this gene was included as part of the GIN analysis, highlighting the importance of integrative analytical approaches that extend beyond simple catalogues of DE genes. The protein product of the *ENPP2* gene interacts with IL-13 during the IL-13-mediated sarcoidosis granulomatous response (Locke *et al*. 2019) and the product of ENPP2 (aka autotaxin) phospholipase action, lysophosphatidic acid, converts recruited monocytes from bone marrow into macrophages (Ray & Rai 2017). Increased levels of PLG (plasminogen) has been shown to encourage granuloma formation; a recent study showed that *M. bovis* BCG bacilli coated in PLG leads to a decrease in phagocytosis, and an increase in granuloma formation (Echeverria-Valencia *et al*. 2019). TNFSF4 has been proposed as an inhibitor to granuloma formation; polymorphisms in the *TNFSF4* gene are associated with Sjogren’s syndrome, phenotypic effects of which include irregular granuloma formation (Nordmark *et al*. 2011; Sun *et al*. 2013). Other human genes that were associated with enriched SNPs include *PLAC8, LDB2, PLOR2F, GNG12*, and *PRR5l*, which have roles in inflammation, pathway initiation and recruitment, but no direct link to granuloma formation for these genes has yet been established.

From the host’s perspective, the ideal outcome of granuloma formation in TB disease is elimination or sterilisation of macrophages infected with pathogenic MTBC strains. However, in cases where the granuloma becomes necrotic, it is clearly beneficial to the pathogen and transmission of the infection (Guirado & Schlesinger 2013). As such, formation of the granuloma can be interpreted as a failure of alveolar macrophages to sufficiently control MTBC pathogens. Consequently, a better understanding of host genes and genomic variation that determines and influences formation and success of the granuloma could lead to novel therapeutics, diagnostics, prognostics, and in the case of livestock, potential targets for genome editing and genome-enabled breeding programmes for bTB disease resistance/resilience (Banos *et al*. 2017; Tsairidou *et al*. 2018; Bishop & Van Eenennaam 2020). The results presented here relating to granuloma formation also emphasise the importance of *M. bovis* infection in cattle and bTB disease as an animal model of hTB (Van Rhijn *et al*. 2008; Waters *et al*. 2014; Buddle *et al*. 2016).

In addition to genes associated with granuloma formation, many of the 32 bovine and human genes identified through bTB and hTB GWAS integration also encode important components of the NF-κB signalling pathway. The nuclear factor-κB (NF-κB) complex constitutes a family of inducible transcription factors that regulate a large array of genes associated with innate and adaptive immunity and inflammatory responses (Zhang *et al*. 2017). Including both bovine and human genes, there were three genes encoding activators (*GNG12, IL17, PELI2)*, two genes encoding inhibitors (*TNFSF4, IER3, SOC3*) and twelve genes encoding downstream targets (*GRM7, CACNA2D3, CSF3, CYB5R3, CXCL13, KRT79, NKG7, PRR5L, SOCS3, SP3, VEGFA, HPSE*). Notable downstream target genes included *HPSE*, which encodes a lung injury serum protein that promotes neutrophil adherence and inflammation (Kedia *et al*. 2018) and *GRM7*, which encodes a protein that modulates adaptive immunity and inflammation (Fallarino *et al*. 2010). Many of the downstream targets are also regulators of granuloma initiation and formation, again highlighting the importance of this feature of TB disease pathogenesis. Because NF-κB-related genes were highlighted in the original GWAS integration, a supplementary analysis was conducted where 17 NF-kB subunit, activator and inhibitor genes (*NFKB1, NFKB2, RELA, RELB, REL, NFKBIA, NFKBIB, NKRF, NFKBIL1, NFKBIZ, NFKBIE, NDFIP1, NKAP, NKAPL, NKIRAS2, NFKBID,* and *NKAPP1*) were directly used as an input gene set for GWAS integration using the *gwinteR* tool (**Supplementary Information File 11**). This resulted in detection of 163 significantly enriched SNPs (6 human and 157 bovine) within or proximal to these NF-κB-associated genes.

It is important to note that the disparity between the human and bovine SNP enrichment for NF-κB genes may be a consequence of the phenotypes used in the bTB GWAS and the hTB GWAS. The bTB phenotype is a susceptibility/resistance sire EBV generated from *M. bovis* infection diagnostic epidemiology data (Ring *et al*. 2019). Conversely, the hTB case-control GWAS phenotype is more directly related to disease resilience because a relatively large proportion of the hTB control cohort (>20%) may have been latently infected with *M. tuberculosis* (Houben & Dodd 2016; Cohen *et al*. 2019). Consequently, the overrepresentation of bovine genes with enriched SNPs in the NF-κB signalling pathway may reflect the importance of innate immune responses in the phenotype used for the bTB GWAS. The hTB case-control GWAS phenotype, on the other hand, may be more directly associated with adaptive immunity (Pai *et al*. 2016; Bloom *et al*. 2017; Simmons *et al*. 2018).

## 5. Conclusion

Although *M. bovis* and *M. tuberculosis* are 99.95% similar at the nucleotide level, they exhibit specific host tropism to sustain across different hosts yet generate comparable disease in their preferred host. In this regard, key to further understanding this host tropism will be elucidation of the shared and species-specific immunological mechanisms underpinning the bovine and human host responses to establishment of infection by *M. bovis* and *M. tuberculosis*, respectively. In addition, identification of critical mycobacterial infection response pathways shared between the two species underlines the importance of the bovine model for understanding human TB. Using four transcriptomics data sets that represent the responses to infection of bAM, hAM, and hMDM, common and distinct genes and gene response pathways across both host species have been identified. Using three different analysis pipelines, a key observation was the role played by the cytokine-cytokine receptor interaction pathway, which was the most enriched pathway in all four experimental groups across seven functional databases. After integration of the downstream functional genomics outputs with the GWAS data sets, 32 bovine and human genes contained or were proximal to SNPs significantly associated with resistance to infection or disease resilience. A striking feature of this result is that 11 of these genes, across all four groups, are directly involved with the formation of the granuloma, while 18 are involved with NF-kB signalling. This indicates that the overall response pathways and gene regulatory networks (GRNs) are comparable across host species and mycobacterial strains. Our work therefore provides underpinning data with which to elucidate the molecular basis of host tropism across the animal- and human-adapted MTBC.

## Data availability

The bovine RNA-seq data set was generated by the authors and can be obtained from the NCBI Gene Expression Omnibus (GEO): accession identifier GSE62506. The human RNA-seq generated by Papp et al can be obtained from the NCBI GEO with accession identifier: GSE114371. Bovine GWAS summary statistics data were obtained from a published study that provides additional information about sequence and genotype data availability (Ring *et al*. 2019). The human GWAS data set for resistance to infection by *M. tuberculosis* used for the integrative genomics work in this study was obtained from the UK Biobank *GeneATLAS* (http://geneatlas.roslin.ed.ac.uk). The computer code required to repeat and reproduce the analyses is available from a GitHub repository (https://github.com/ThomasHall1688/Human_Bovine_comparison). The computer code used to integrate the functional genomics outputs and GWAS data sets is available from a GitHub repository (https://github.com/ThomasHall1688/gwinteR) Bovine RNA-seq filtering and mapping statistics are available from the Dryad Digital Repository: https://doi.org/10.5061/dryad.83bk3j9q6.

## Author contributions

TJH, SVG and DEM conceived and designed the study. JAB performed experimental work. SCR and DPB provided cattle genotype and other genomic data. TJH, GJM, MPM, KEK, and JAW performed bioinformatics and computational analyses. TJH and DEM wrote and prepared the manuscript and figures. All authors read and approved the final manuscript.

## Acknowledgements

The authors would like to thank Donagh Berry and Siobhán Ring for provision of cattle GWAS data. We would also like to thank Carol Correia, Joseph Crispell, Eamonn Gormley, and Kerri Malone for useful discussions.

## Supplementary information

Supplementary information for this article can be found online.

