## Supplementary Material for "Integrative and comparative genomic analyses of mammalian macrophage responses to intracellular mycobacterial pathogens"

[c] Irish Cattle Breeding Federation, Bandon, Cork, P72 X050, Ireland.

[d] Teagasc, Animal and Grassland Research and Innovation Centre, Moorepark, Cork, P61 C996, Ireland.

[e] UCD School of Veterinary Medicine, University College Dublin, Belfield, Dublin, D04 V1W8, Ireland.

[f] UCD Conway Institute of Biomolecular and Biomedical Research, University College Dublin, Belfield, Dublin, D04 V1W8, Ireland.

[1] Current address: Novartis Ireland Ltd., Elm Park Business Campus, Dublin, D04A9N6, Ireland.

[*] **Corresponding author:**
David E. MacHugh
Animal Genomics Laboratory, UCD School of Agriculture and Food Science, UCD College of Health and Agricultural Sciences, University College Dublin, Belfield, Dublin D04 V1W8, Ireland.

### Supplementary Figures

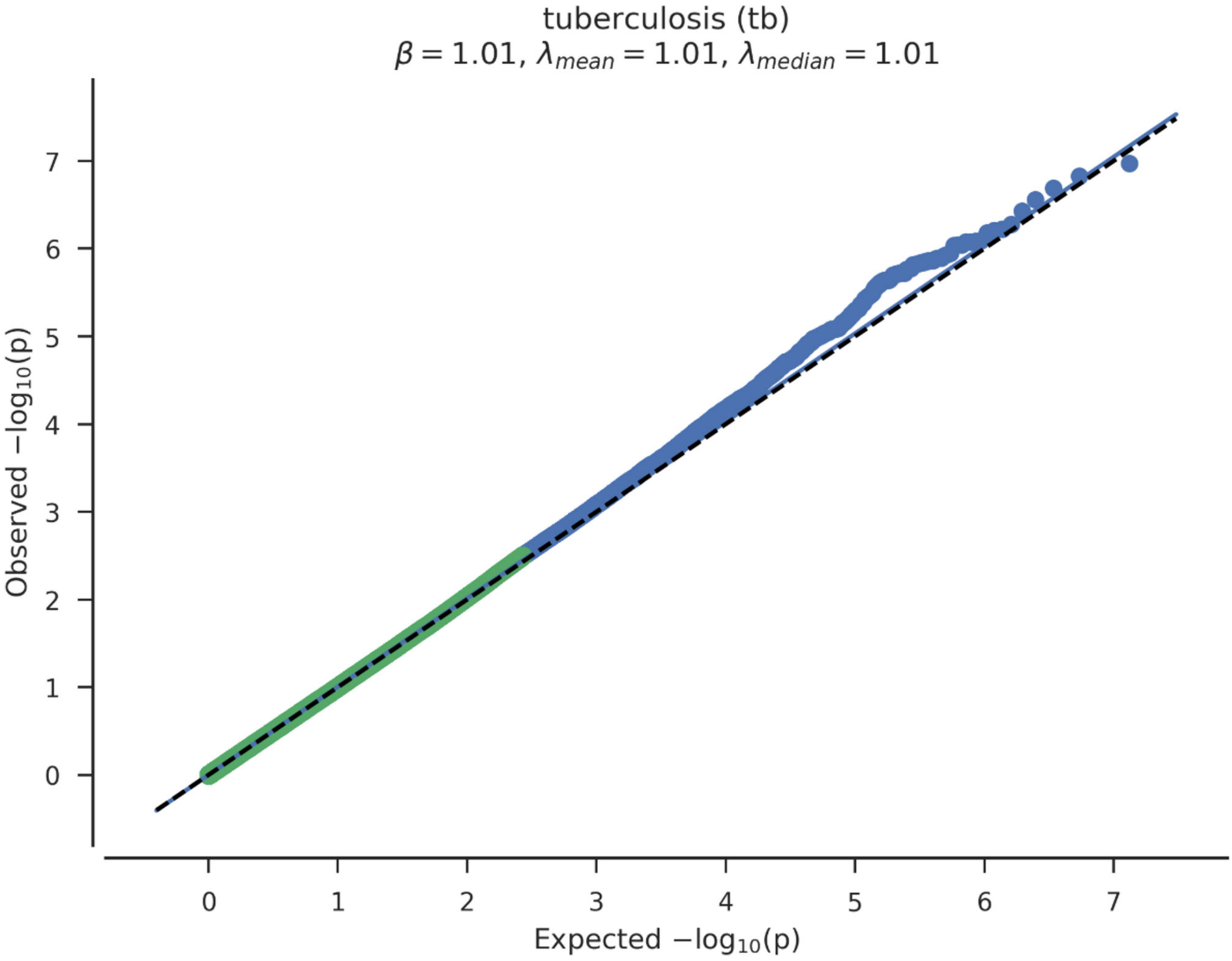


**Supplementary Figure 1:** Human TB GWAS SNP Q-Q Plot containing 9,113,133 imputed variants that passed quality control (modified from http://geneatlas.roslin.ed.ac.uk/trait/?traits=190).

A)

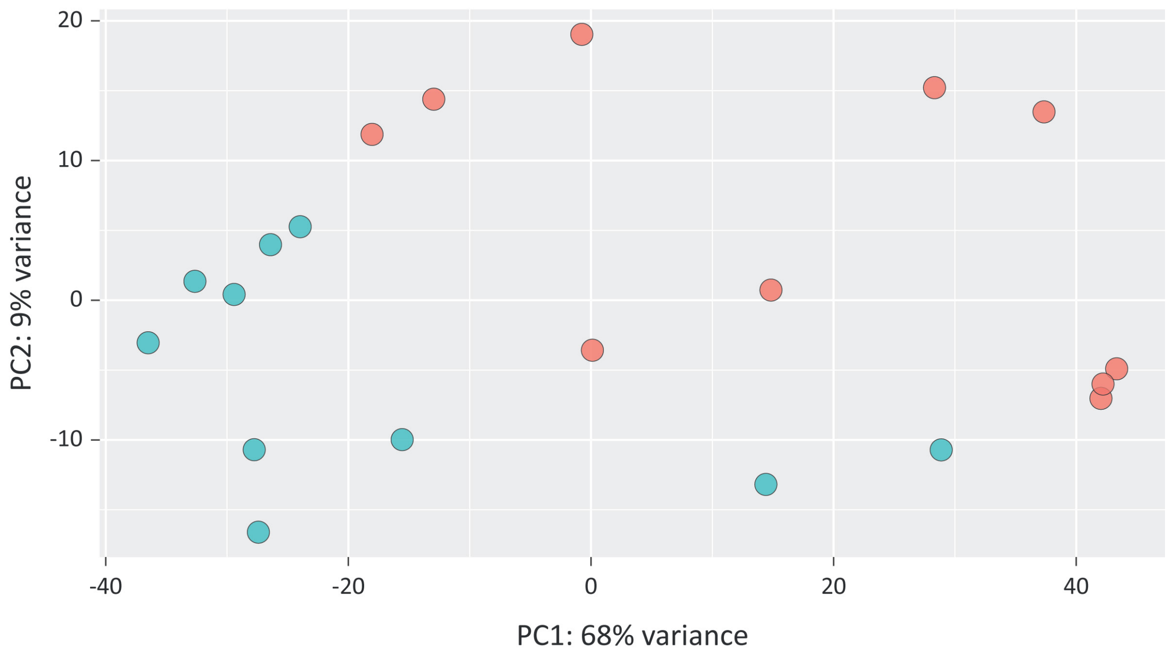


B)

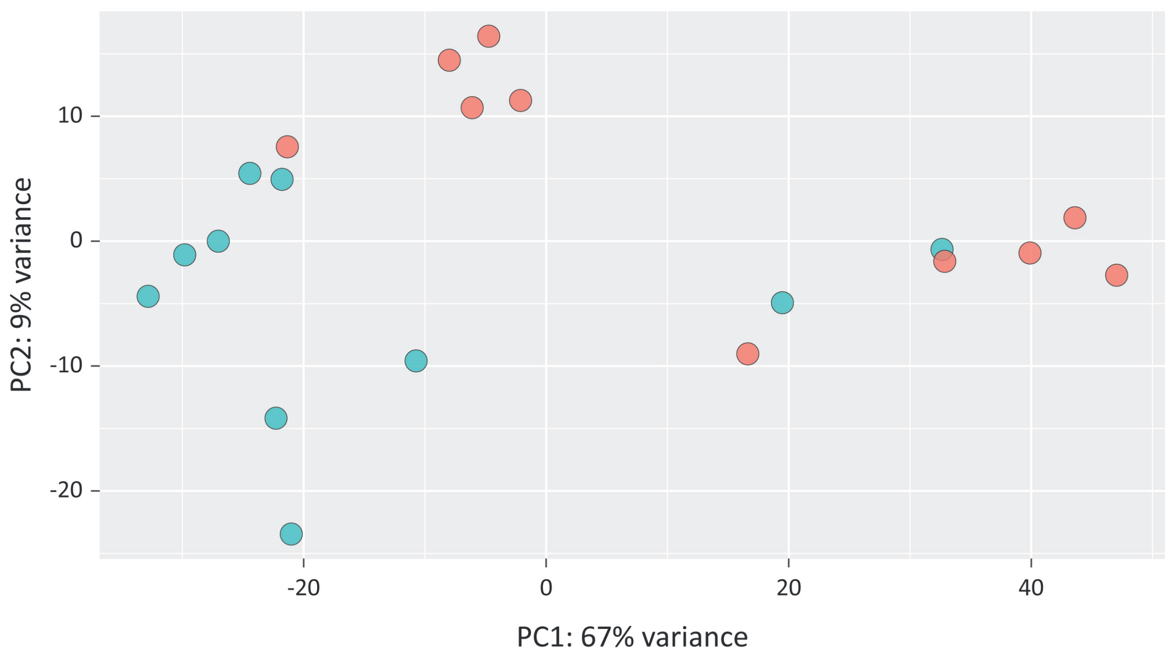


**Supplementary Figure 2**: Principal component analysis (PCA) plots for individual animal bAM gene expression at 24 hpi. **A.** *M. bovis*-infected, and **B.** *M. tuberculosis*-infected. Red indicates infected animals and blue indicates non-infected control animals.

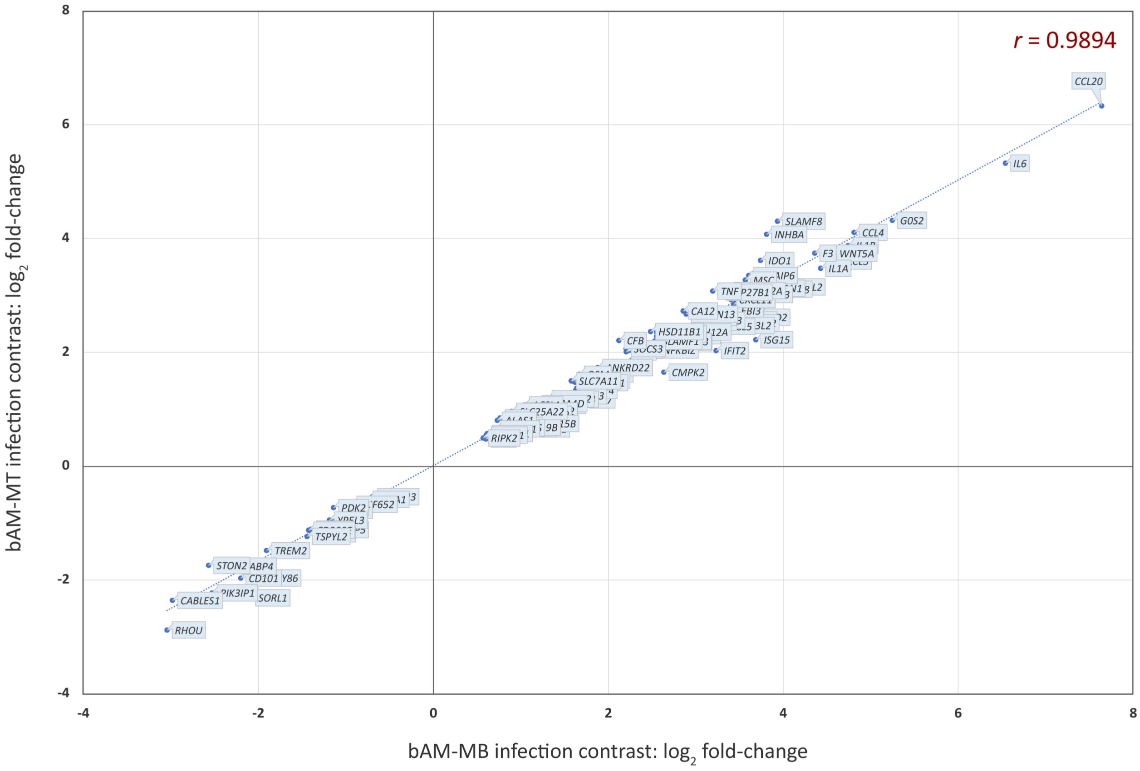


**Supplementary Figure 3**: Correlation plot of the 118 DE genes common across all experimental groups at 24 hpi for the *M. tuberculosis*-infected bAM versus the *M. bovis*-infected bAM.

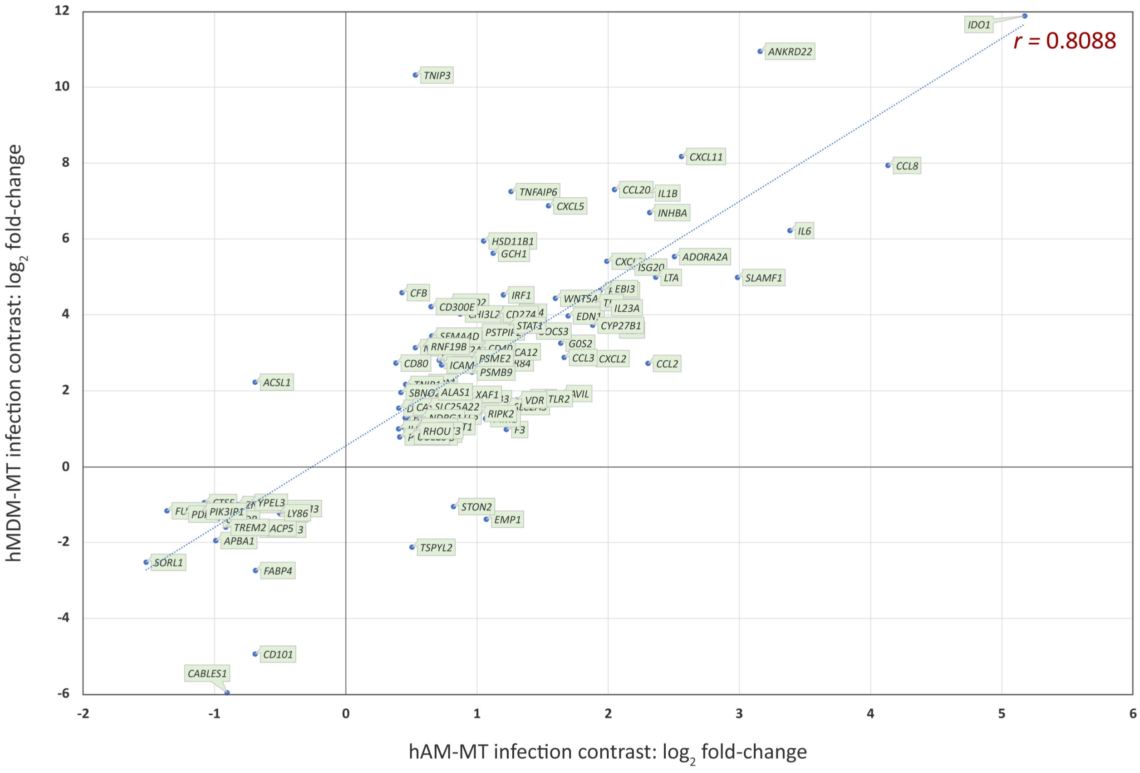


**Supplementary Figure 4**: Correlation plot of the 118 DE genes common across all experimental groups at 24 hpi for the *M. tuberculosis*-infected hMDM versus the *M.* tuberculosis-infected hAM.

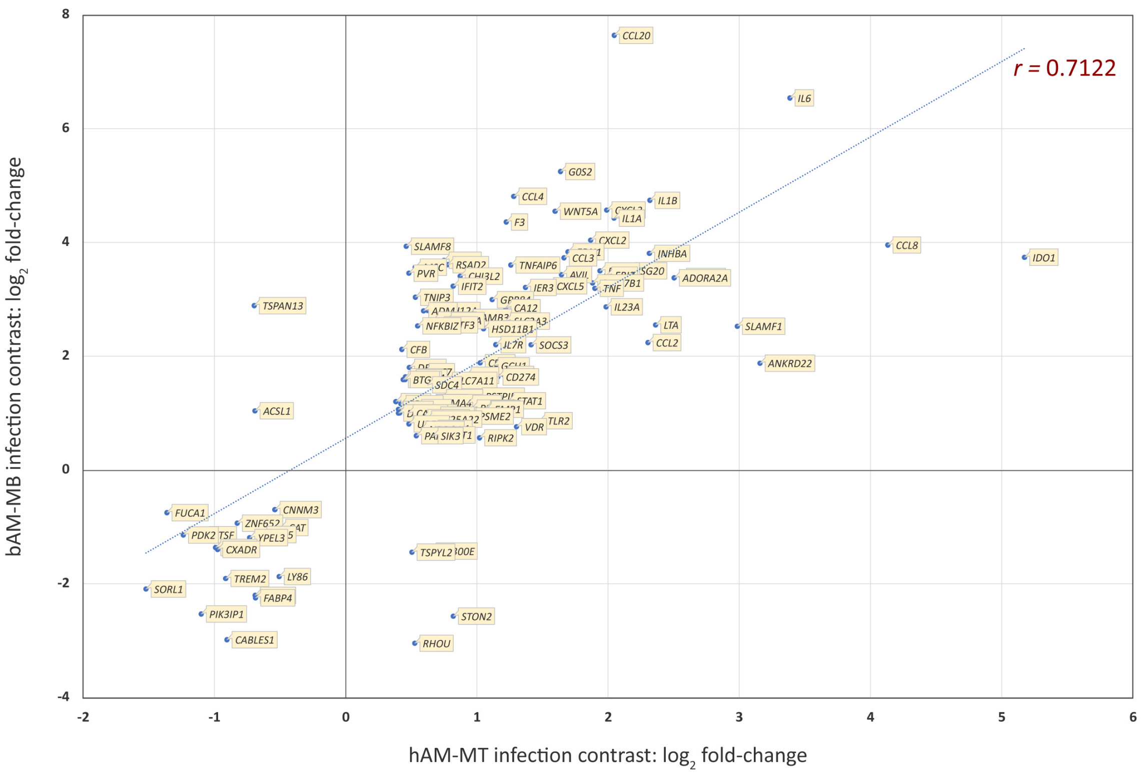


**Supplementary Figure 5**: Correlation plot of the 118 DE genes common across all experimental groups at 24 hpi for the *M. bovis*-infected bAM versus the *M. tuberculosis*-infected hAM.

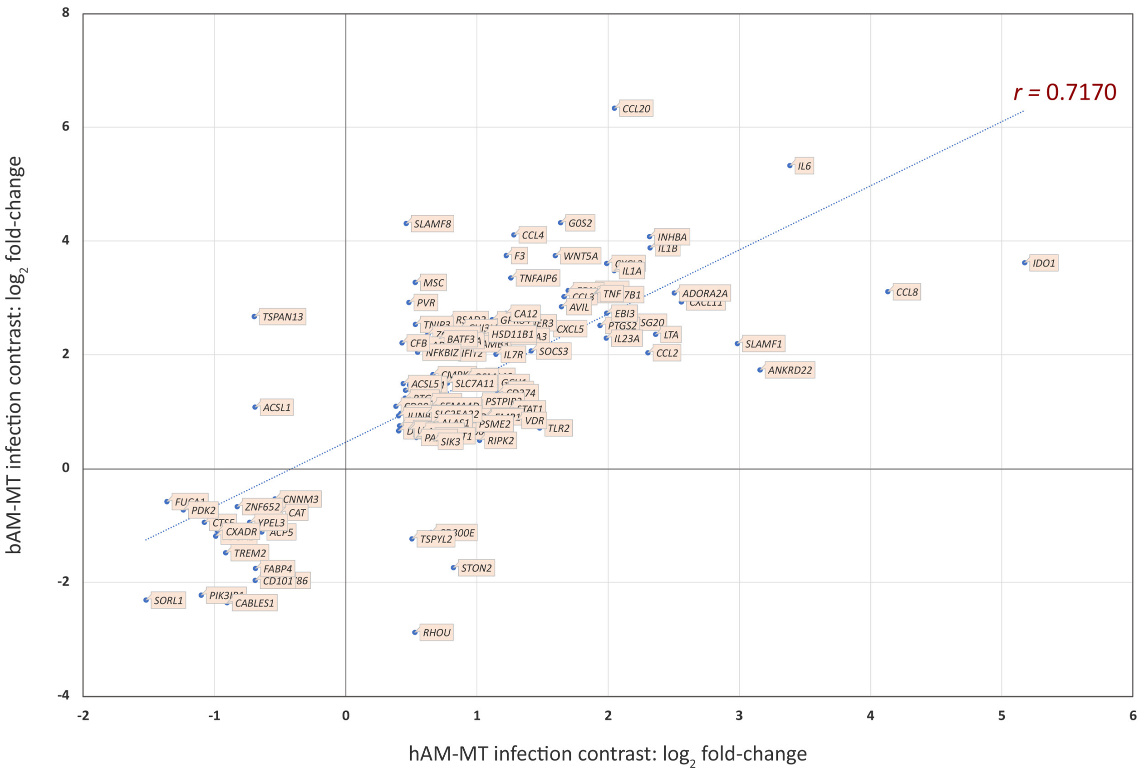


**Supplementary Figure 6**: Correlation plot of the 118 DE genes common across all experimental groups at 24 hpi for the *M. tuberculosis*-infected bAM versus the *M. tuberculosis*-infected hAM.

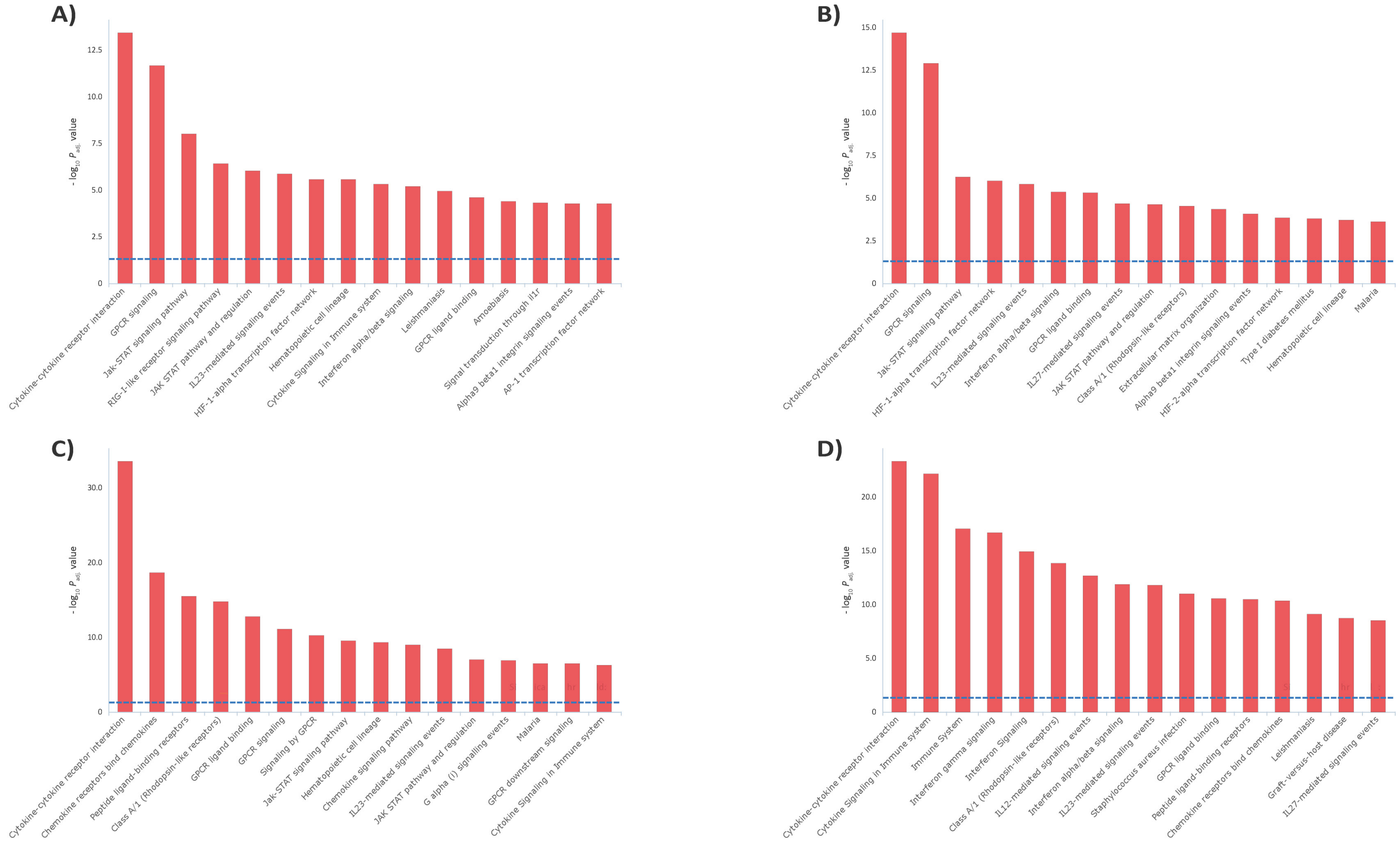


**Supplementary Figure 7: Top 16 enriched biological pathways for each experimental group from the combined pathway analysis (CPA). A.** *M. bovis* -infected bAM. **B.** *M. tuberculosis*-infected bAM. **C.** *M. tuberculosis*-infected hAM. **D.** *M. tuberculosis*-infected hMDM. The 0.05 $P_{adj.}$ threshold is represented by a blue dashed line. All pathways shown were upregulated.
